# Live-cell microscopy or fluorescence anisotropy with budded baculoviruses - which way to go with measuring ligand binding to M_4_ muscarinic receptors?

**DOI:** 10.1101/2021.12.22.473643

**Authors:** Maris-Johanna Tahk, Jane Torp, Mohammed A.S. Ali, Dmytro Fishman, Leopold Parts, Lukas Grätz, Christoph Müller, Max Keller, Santa Veiksina, Tõnis Laasfeld, Ago Rinken

**Author notes:** Corresponding author, E-mail addresses. Maris-Johanna Tahk and Jane Torp contributed to this article equally.

## Abstract

M_4_ muscarinic receptor is a G protein-coupled receptor that has been associated with alcohol and cocaine abuse, Alzheimer’s disease and schizophrenia which makes it an interesting drug target. For many G protein-coupled receptors, the development of high-affinity fluorescence ligands has expanded the options for high throughput screening of drug candidates and serve as useful tools in fundamental receptor research. So far, the lack of suitable fluorescence ligands has limited studying M_4_ receptor ligand binding. Here, we explored the possibilities of using fluorescence-based methods for studying binding affinity and kinetics to M_4_ receptor of both labeled and unlabeled ligands. We used two TAMRA-labeled fluorescence ligands, UR-MK342 and UR-CG072, for assay development. Using budded baculovirus particles as M_4_ receptor preparation and fluorescence anisotropy method, we determined the affinities and binding kinetics of both fluorescence ligands. The fluorescence ligands could also be used as reported probes for determining binding affinities of a set of unlabeled ligands. Based on these results, we took a step further towards a more natural signaling system and developed a method using live CHO-K1-hM_4_R cells and automated fluorescence microscopy suitable for routine determination of unlabeled ligand affinities. For quantitative image analysis, we developed random forest and deep learning-based pipelines for cell segmentation. The pipelines were integrated into the user-friendly open-source Aparecium software. Both developed methods were suitable for measuring fluorescence ligand saturation binding, association and dissociation kinetics as well as for screening binding affinities of unlabeled ligands.

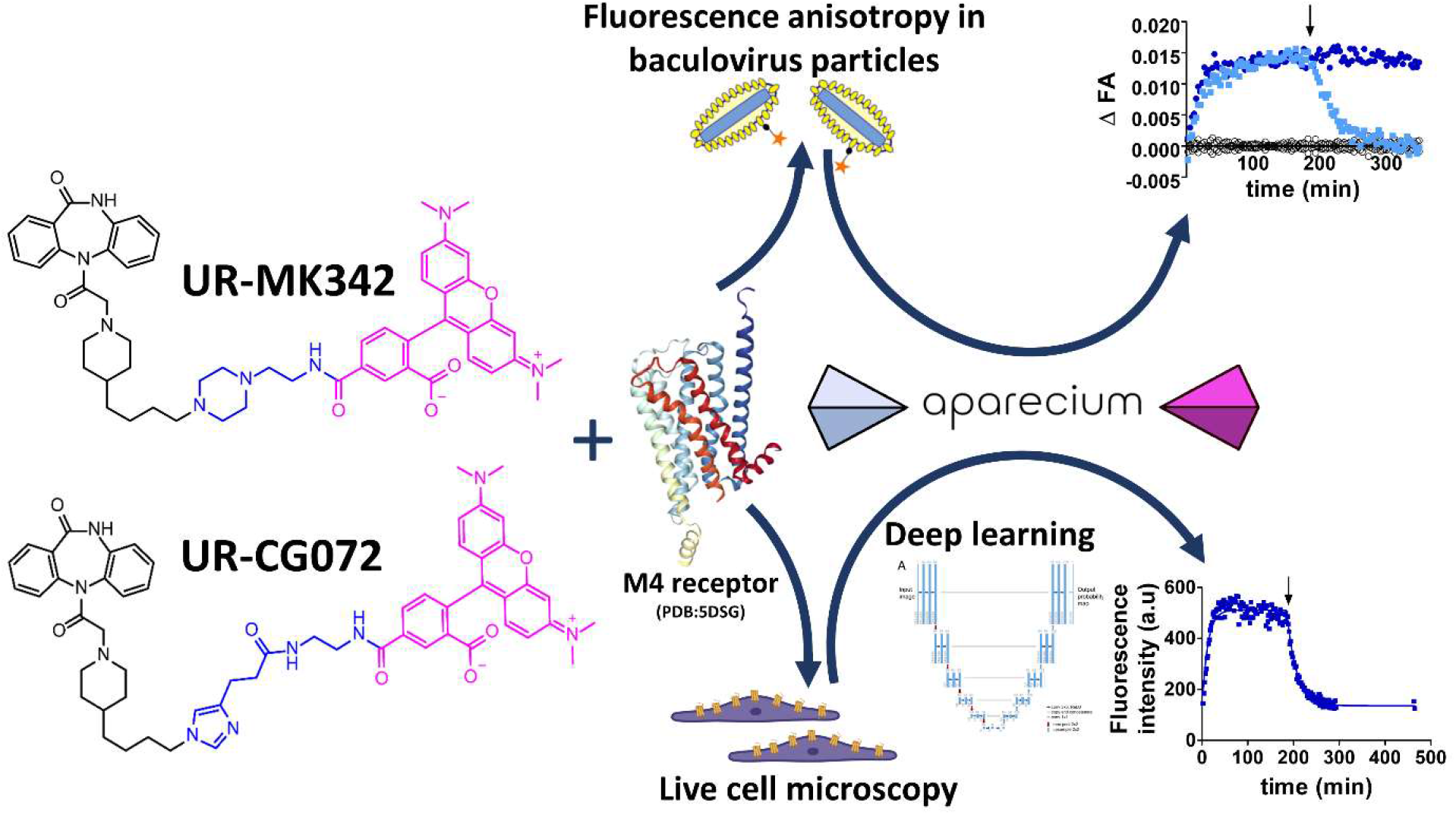

## 1. Introduction

Muscarinic acetylcholine receptors (mAChR) are a group of G-protein coupled receptors (GPCRs) with five subtypes M_1_-M_5_ which play a crucial role, e.g. in the regulation of memory, heart and bladder function and dopaminergic neurotransmission [1–4]. Lately, the M_4_R was suggested to be a potential drug target for the treatment of neurodegenerative and neuropsychiatric disorders like Alzheimer’s disease or schizophrenia [5, 6]. Furthermore, as new links emerge between the M_4_ receptor and alcoholism, as well as a between M_4_ receptor polymorphism and cocaine and heroin abuse, the M_4_ receptor becomes an even more versatile drug target [7, 8]. Despite the growing importance, the development of novel drugs targeting the M_4_ receptor is difficult as the similarity of orthosteric binding sites of all mAChR leads to low subtype selectivity of ligands [9]. One solution is the development of allosteric modulators, which may exhibit higher subtype selectivity but relatively lower affinities [10]. To find suitable drugs, ligand screening remains an important step in the drug development process. Screening for new drug candidates using fluorescence methods has become quite popular due to several advantages over radioligand based assays [11]. However, until now only a limited number of fluorescence ligands have been available for mAChR and to our knowledge none have been extensively used to develop assays to study M_4_ receptors [12–14]. Recently, several novel low molecular weight fluorescently labeled ligands targeting mAChRs were described [15, 16]. Of these ligands, TAMRA labeled UR-CG072 and UR-MK342 have already been successfully used for studying M_2_ receptors in nanoBRET and fluorescence anisotropy (FA) assays [17]. Even though these ligands show a slight preference for the M_2_ receptor, they still have high affinity for M_1_ and M_4_ receptors. Therefore, the new fluorescent ligands should be suitable as probes for studying M_4_ receptors in drug candidate screening assays as well as in a large variety of fluorescence microscopy techniques from live tissue systems to single-molecule studies [18–20].

One of the most common options for characterizing fluorescent probe binding to proteins including GPCRs is the FA method [21–25]. For the successful development of FA assays, several unique aspects must be considered. Most importantly, FA is a ratiometric assay with its value depending on the ratio of bound and free ligand. Therefore, all experiments must be designed in a way that the probe and receptor concentrations are in a similar range, which means that ligand and receptor depletion should be taken into account [26]. The main advantage of the FA method is that there is no need to separate bound ligand from free ligand, making it easy to continuously collect time-course data during ligand binding. These time course data can be used to obtain kinetic parameters and to develop reaction kinetics models of ligand binding for more insight into the complicated regulation of signal transduction. In addition to cell membranes, budded baculovirus (BBV) particles can serve as a high-quality receptor source for FA assays. BBV particles are advantageous because they have a fixed shape (∼50 nm x 300 nm) and homogeneous size distribution, resulting in minimal noise and small variability between replicates compared to membrane preparations [26–28]. Due to the small size and low sedimentation rate of BBV particles they are well suited for performing homogeneous assays. However, downstream signaling cascades are not present BBV particles. Furthermore, BBV particles are produced in Sf9 insect cells, where the membrane composition differs from mammalian systems. Most of these problems can be avoided by using more natural live-cell assays for receptor display. Among multiple developed assays [29], NanoBRET has gained a lot of popularity in recent years due to its homogeneous format, possibility of real-time measurements and relatively good compatibility with a wide array of fluorophores. However, it requires genetically modified receptors, which may have an influence on ligand binding and receptor activation [30]. Studying wild type receptors is more difficult, as the receptor cannot be tagged which in turn does not allow to take advantage of the high sensitivity of bioluminescence approaches. Further, the plate reader based RET methods only provide cell population average statistics instead of single-cell resolution information which may hide some important effects. One solution to both problems is flow cytometry, which can measure fluorescent ligand binding to individual cells. However, it cannot follow binding to a single cell over time nor spatially resolve from which part of the cell the fluorescence originates from [31]. In contrast, high throughput microscopy can provide spatial information as well as time-course information for the same cells, making more detailed analysis possible. On the downside, extracting pharmacologically relevant quantitative information from the bioimages requires more complex data analysis algorithms. However, once an automated data analysis solution with user-friendly software exists, it can be re-used in future studies.

Microscopy methods open many possibilities for assay setup, but performing time-resolved measurements with the cellular resolution is not trivial and existing methods have several potential issues [32]. In previously published works kinetics of ligand binding to live cells in an HTS compatible manner have only been analyzed by fluorescence intensity of the whole image [33, 34]. For these methods, it is necessary to seed cells consistently as a high confluency monolayer, but this is either difficult or practically impossible to achieve with some cell lines [35]. Furthermore, it is much more difficult to identify individual cells from a dense monolayer thus reducing the number of parameters that can be studied. In addition, dense monolayers can significantly affect physicochemical environmental parameters such as oxygen concentration which can also have more direct effects on muscarinic receptor signaling [36]. For example, transient hypoxic conditions lead to increased phosphorylation of M_1_ and M_2_ receptors [37]. Finally, dense cell monolayers can easily cause focusing errors in automated microscopy, as some cells may have detached or formed a second layer. A better approach was developed with HEK-293-D_3_R cells, which uses a machine-learning algorithm for detecting only the fluorescence intensity originating from cell membranes in equilibrium conditions and does not rely on dense monolayers [35]. However, ligand binding kinetics were not analysed in that study. Nevertheless, kinetic measurements should be possible with a similar setup after adjusting the experimental design and the image analysis pipeline.

The most difficult steps of microscopy image analysis are usually cell detection and segmentation, which is necessary for robust quantification of the fluorescence signal. Approaches for these tasks have gone through a paradigm shift from classical computer vision techniques to machine learning and especially deep learning (DL) methods. Deep neural networks dominate most of the developed benchmark datasets for general problems as well as bioimage analysis specifically [38–40]. A large number of DL architectures have been developed over the past few years, but their wide application can be limited by compatibility issues with popular image analysis software and too complex design for comprehensive understanding for life-scientists [41–48]. Therefore, a widely supported and well known U-Net architecture is used in the present study for cell segmentation from bright-field images, as it has shown good results for similar microscopy images [43, 49].

In this study, we developed new fluorescence-based ligand binding assays for the M_4_ receptor. These assays utilize two recently developed 5-TAMRA labeled dibenzodiazepinone derivatives, UR-MK342 and UR-CG072 [15], and two different receptor sources. As both BBV particle-based FA and live cell based microscopy assays have distinct advantages, we studied and compared the two options and discovered, that both options are viable. To our best knowledge, this is the first detailed description of M_4_ receptor fluorescence ligand binding assays, which opens up many new possibilities to study these receptors.

## 2. Materials and methods

### Materials

Assay buffer consisted of MilliQ water, 135 mM NaCl (AppliChem, Darmstadt, Germany), 1 mM CaCl_2_ (AppliChem), 5 mM KCl (AppliChem), 1 mM MgCl_2_ (AppliChem), 11 mM Na-HEPES (pH = 7.4) (Sigma-Aldrich, Taufkirchen, Germany), protease inhibitor cocktail (according to the manufacturer’s description, Roche, Basel, Switzerland) and 0,1% Pluronic^®^ F-127 (Sigma-Aldrich).

Muscarinic acetylcholine receptor ligands acetylcholine, arecoline, pirenzepine, pilocarpine, atropine and scopolamine were purchased from Sigma-Aldrich and carbachol from Tocris Bioscience (Abingdon, United Kingdom). The syntheses of the fluorescent ligands UR-MK342 and UR-CG072 [15] and the dualsteric M_2_R ligands UR-SK59[50], UR-SK75 [50] and UNSW-MK259 [51], showing also high M_4_R affinity, were described previously. All ligand stock solutions were prepared using cell culture grade DMSO (AppliChem) and stored at -20 °C.

### Cell culture

*Spodoptera frugiperda* Sf9 (Invitrogen Life Technologies, Schwerte, Germany) cells were maintained as a suspension culture in serum-free insect cell growth medium EX-CELL 420 (Sigma-Aldrich) at 27 °C in a non-humidified environment.

Non-transfected Chinese hamster ovary cells (CHO-K1) were purchased from ATCC, LGC Standards (Wesel, Germany) and CHO-K1 expressing human M_4_ receptor (CHO-K1-hM_4_R cells) were obtained from Missouri S&T cDNA Resource Centre (Bloomsberg, USA). CHO-K1 cells were cultured in DMEM/F12 (Sigma-Aldrich) with 9% FBS (Sigma-Aldrich), antibiotic antimycotic solution (100 U/ml penicillin, 0,1 mg/ml streptomycin, 0,25 μg/ml amphotericin B) (Sigma-Aldrich) and for CHO-K1-hM_4_R 750 μg/mL of selection antibiotic geneticin (G418) (Capricorn Scientific, Ebsdorfergrund, Germany) were added. All mammalian cells were grown in a humidified incubator at 37 °C with 5% CO_2_ until 90% confluency. To detach the cells from the plate 0.05% trypsin with EDTA (Gibco, Paisley, Scotland) was used.

Cell culture viability and density were determined with an Automated Cell Counter TC20™ (Bio-Rad Laboratories, Sundyberg, Sweden) by the addition of 0.2% trypan blue (Sigma-Aldrich). All experiments with CHO-K1-hM_4_R, CHO-K1 and Sf9 cell cultures were performed with passages 40-50, 33 and 2-25 respectively. Mammalian cell-lines were tested and determined to be mycoplasma-negative.

### Preparation of budded baculovirus particles

The human M_4_ receptor in pcDNA3.1+ was purchased from cDNA Resource Center (www.cdna.org) and manufacturing and production of BBV containing human M_4_ receptor were performed as described in [23] with some modification. For cloning M_4_ into pFastBac vector, BamHI and XbaI sites were used with enzymes from (Thermo Fisher Scientific (Schwerte, Germany). To transform the bacmid into Sf9 cells the transfection reagent FuGene 6 (Promega Corporation, Madison, USA) was used according to the manufacturer’s protocol. After the viruses were generated and collected, the amount of infectious viral particles per ml (IVP/ml) for all the baculoviruses was determined with the Image-based Cell Size Estimation (ICSE) assay [52].

To produce the BBV particles, Sf9 cells were infected with MOI = 3 and incubated for 4 days (end viability of Sf9 cells was 55%). The supernatant, containing BBV particles, was gathered by centrifugation for 15 min at 1600 g. Next, the BBV particles were concentrated 40-fold by high-speed centrifugation (48000 g at 4 °C) for 40 min followed by washing with the assay buffer and homogenization with a syringe and a 30G needle. The suspension was divided into aliquots and stored at -90 °C until the experiments. BBV particle preparations were done several times. Receptor concentration for the BBV particle stocks was estimated R_stock_UR-CG072_= 9.7 ± 1.1 nM and R_stock_UR-MK432_= 5.5 ± 0.7 nM, using the model described in [28].

### Fluorescence anisotropy experiments

FA experiments were carried out on black flat bottom half-area 96 well plates (Corning, Glendale, USA). A suitable combination of the fluorescent ligand, competitive ligand and BBV particle suspension was added to each well. Assay buffer was added so that the final liquid volume in each well was 100 µL.

In saturation binding experiments, two concentrations of fluorescent ligands were used, 2 nM and 20 nM for UR-CG072 and 1 nM and 6 nM for UR-MK342. 2 µM or 20 μM UNSW-MK259 in the case of UR-CG072 and 1 µM or 6 μM scopolamine in the case of UR-MK342 were used for determination of nonspecific binding. Two-fold serial dilutions of BBV particle suspension was added starting from 60 µL. Wells without BBV particles were used as a free fluorescent ligand control.

For competition binding experiments the concentrations of fluorescent ligands UR-CG072 and UR-MK342 were kept constant at 5 nM and the volume of BBV particles was also kept constant at 20 μL (C_final_ ≈ 1 - 2.2 nM). Five- or six-fold serial dilutions of the competitive ligands were used. Also, replicate wells with no competitive ligand were included and for blank correction replicate wells with only BBV particles was included. Measurements were carried out with 3 min intervals for 13-15 h at 27 °C. A custom made glass lid was used in all the experiments to minimize the evaporation from the wells. In all cases, BBV particles were added as the last component to initiate the ligand binding process.

For kinetic experiments, 5 nM UR-CG072 or 6 nM UR-MK342 was used. In nonspecific binding wells, 6 µM or 3 µM scopolamine was added, respectively. The reaction was initiated by addition of 20 µL of M_4_ receptor displaying BBV particles. After 180 min the dissociation was initiated by the addition of 2 µL of 300 µM (C_final_ = 6 µM) or 150 µM (C_final_ = 3 µM) scopolamine for UR-CG072 or UR-MK342, respectively. 2 µL of assay buffer was added instead of the competitive ligand to association kinetics wells to maintain the equivalent volume in all wells.

In all experiments, the fluorescence intensity values were blank corrected for BBV particle autofluorescence by subtracting the respective parallel or perpendicular fluorescence intensity value of a blank well from the respective measurement well. The blank wells contained no ligands but only the same concentration of BBV particles as the measurement well.

FA measurements were performed with multi-mode plate reader Synergy NEO (BioTek Instruments, Winooski, USA), which is equipped with a polarizing 530(25) nm excitation filter and 590(35) nm emission filter allowing simultaneous parallelly and perpendicularly polarized fluorescence detection. At least three individual experiments were carried out in duplicate.

### Microscopy of DiI stained CHO-K1-hM4R cells

CHO-K1-hM_4_R cells were grown as described above and seeded with a density of 25 000 cells/well into a µ-Plate 96 well Black plate (Ibidi, Gräfelfing, Germany) 5 hours before the experiment. A stock solution of 1 mM DiI (Invitrogen, Eugene, Oregon, USA) in DMSO stored at -20 °C was thawed and sonicated in an ultrasound bath for 5 min to disrupt aggregates. Cell medium was removed and replaced with 200 µL/well of 2 µM DiI in DPBS with Mg^2+^ and Ca^2+^ (Sigma-Aldrich) to stain the cell membranes. The cells were incubated with the solution for 10 minutes before imaging. The cells were imaged with Cytation 5 cell imaging multi-mode plate reader equipped with 20X LUCPLFLN objective (Olympus) from Bright-field and RFP channels (LED light source with excitation filter 531(40) nm and emission filter 593(40) nm for RFP channel (BioTek Instruments) with the following parameters for bright-field: LED intensity = 4, integration time = 110 ms, camera gain = 24 and for RFP fluorescence channel: LED intensity = 1, integration time = 71 ms, camera gain = 24. The cells were imaged in the montage mode (196 locations) with Z-stack (10 planes, 4 planes below and 5 planes above focus) to cover any imaging location-dependent variability and simulate potential autofocusing errors.

### Live-cell ligand binding imaging

CHO-K1-hM_4_R cells were seeded into µ-Plate 96 well Black plate (Ibidi) at densities of 25 000 - 30 000 cells/well in DMEM/F-12 medium and incubated for 5-7 h. Immediately before the measurement, the cell culture media was exchanged for the same cell culture media containing ligands. At all times, the well volume was kept at 200 µL.

For determining UR-CG072 affinity to the M_4_ receptor, saturation binding experiments were carried out using two-fold dilutions of UR-CG072 starting from 8 nM. Nonspecific binding was measured in the presence of 3.7 μM scopolamine. The cells were incubated with ligands in Cytation 5 at 5% CO_2_ and 37 °C for 2 h before imaging.

For measuring UR-CG072 binding kinetics to M_4_ receptor, 2 nM UR-CG072 was added to the cells and imaging was immediately initiated. To achieve sufficient temporal resolution, only two wells were imaged in parallel. After approximately 3 h of association, 10 µL of 100 µM scopolamine (C_final_ = 5 µM) was added to start dissociation.

The competition binding assay was performed using 2 nM UR-CG072. The different competitive ligand concentrations were pipetted to the plate in randomized order, to avoid correlation between well imaging order and concentration. It was determined that 2 h was sufficient to reach equilibrium for IC_50_ value measurement as the IC_50_ values for scopolamine and carbachol at 2 h and 5 h remained constant within uncertainty limits.

The imaging was performed with Cytation 5 as described above. Saturation binding experiments were performed with following imaging parameters in bright-field: LED intensity = 4, integration time = 110 ms, camera gain = 24 and in RFP fluorescence channel LED intensity = 1 or 2, integration time = 827 ms, camera gain = 24. For kinetic binding assays all the parameters were the same except for RFP fluorescence channel LED intensity = 5. For competition binding assays the imaging parameters used in bright-field were: LED intensity = 5, integration time = 1222 ms, camera gain = 0 and in RFP fluorescence channel: LED intensity = 5, integration time = 613 ms, camera gain = 24 or the same as for kinetic experiments. The cells were imaged in the montage mode (4 locations/well) with Z-stack (10 planes, 4 planes below focal plane, 1 in focus and 5 planes above focal plane).

### Cell segmentation with ilastik software

To develop a bright-field cell segmentation model based on the random forest (RF) algorithm, a total of three ilastik [53] pixel classification models were trained (**RF-FL-1**, **RF-BF-1** and **RF-BF-2**). Two of the models (**RF-FL-1** and **RF-BF-1**) were intermediate helper models used for training the final **RF-BF-2** model. Here, the models are named by combining the model type (RF or U-Net3), an input imaging modality that the model used for cell detection (BF for bright-field images and FL for fluorescence images) followed by the index of the model of the particular type. For developing the **RF-FL-1** model, a set of fluorescence images of CHO-K1-hM_4_R cells stained with fluorescent lipophilic dye DiI was generated. 30 of these images were randomly chosen from different locations of the well for the training set. The images were in-focus (10 images), 3 μm above (10 images) and below (10 images) the focal plane to increase the model robustness against focusing errors. The Gaussian Smoothing, Laplacian of Gaussian, Gaussian Gradient Magnitude, Difference of Gaussians, Structure Tensor Eigenvalues, and Hessian of Gaussian Eigenvalues features were selected for sigma values of 0.70, 1.00, 1.60, 3.50, 5.00, 10.00, 15.00 and 20.00 px. In addition, the Gaussian Smoothing feature with a sigma value of 0.30 px was selected in the ilastik feature selection stage. **RF-FL-1** was set up to perform binary pixel classification using cell and background classes. Some pixels of cells and background were manually annotated by adding annotations over the respective pixels of the in-focus images. More annotations were added at the fringe of cells to enhance the accuracy of the predictions. The annotations of the in-focus images were transferred to the respective out-of-focus images from the same field of view. With these annotations, the random forest (RF) based model (**RF-FL-1**) was trained. Model export was set to generate simple binary segmentation. Then, the cells on the rest of the fluorescence images (186 images) were segmented in the batch processing mode creating a set of masks for 196 fields of view with the ten fields of view remaining in the training set. Next, the binary segmentation images were automatically reclassified into three classes: intracellular area (IC), membrane (MB) and near-membrane background (NMBG) with the rest of the pixels representing background (BG). IC class was generated by image erosion of the predicted cell masks by a 2-pixel radius disk structuring element. MB class was generated by image dilation of IC masks with a 3-pixel radius disk structuring element and pixels were assigned to the NMBG class by further image dilation of the MB images with a 7-pixel radius disk structuring element and excluding pixels already assigned to MB or IC classes. Next, a class balancing step was performed to obtain an equal number of pixels for each of the classes. For that, all of the pixels from the class with the smallest number of pixels (MB) were selected and an equal number of pixels were selected randomly from IC and NMBG classes. The operation was performed for each image separately. Images generated by this process were considered as the ground truth for training **RF-BF-1** model. **RF-BF-1** was trained to detect cells from contrasted projections of bright-field Z-stacks. The Z-stack of bright-field images was converted into a single higher contrast image as described in [35]. Twenty fields of view were used as a training set in **RF-BF-1** for the detection of IC, MB, NMBG areas from the contrast-enhanced bright-field image projections. The Gaussian Smoothing, Laplacian of Gaussian, Gaussian Gradient Magnitude, Difference of Gaussians, Structure Tensor Eigenvalues, and Hessian of Gaussian Eigenvalues features were selected for sigma values of 0.70, 1.00, 1.60, 3.50, 5.00, 10.00, 15.00, 20.00, 25.00, 30.00 and 35.00 px. In addition, the Gaussian Smoothing filter with a sigma value of 0.30 px was selected in the ilastik feature selection stage. Twenty ground truth images generated by **RF-FL-1** were used as labels of IC, MB, NMBG classes in the respective images to train a model for the detection of three classes of pixels from the contrast-enhanced bright-field images. The prediction quality was estimated by the recall, precision, F_1_ score and Matthews Correlation Coefficient (MCC) metrics as shown in Table 1. Classification quality metrics were measured by considering the IC pixels to form a positive class while all other classes (MB, NMBG, BG) were merged to form the negative class. Thus, misclassifications of pixels between MB, NMBG and BG classes had no impact on the quality metrics.

**Table 1.** Quality metrics of the developed machine learning models.

| **Model** | **U-Net3-FL-1** | **RF-FL-1** | **U-Net3-BF-1** | **RF-BF-1** | **RF-BF-2** |
| --- | --- | --- | --- | --- | --- |
| **Metric** |  |  |  |  |  |
| **Recall** | 0.94 | 0.91 | 0.86 | 0.75 | 0.72 |
| **Precision** | 0.88 | 0.94 | 0.93 | 0.77 | 0.74 |
| **F1 score** | 0.91 | 0.93 | 0.89 | 0.76 | 0.73 |
| **MCC** | 0.89 | 0.91 | 0.86 | 0.71 | 0.67 |

Finally, the **RF-BF-2** model was trained to improve the prediction quality of the **RF-BF-1** model by adjusting the class balance by adding ground truth annotation to pixels which the **RF-BF-1** model had failed to classify. The same training set of 20 contrast-enhanced bright-field images was used for the **RF-BF-1** model. In this training run, the fourth class of pixels was created for background (BG) from all the previously unclassified pixels. To create ground truth images for **RF-BF-2**, the class balancing step was performed again as previously described. Additionally, the labels were improved by manually adding pixels to each class, which the **RF-BF-1** model had failed to classify. The same image features were used as in the **RF-BF-1** model. The prediction quality of **RF-BF-2** was evaluated with the test set (Table 1, Fig 7 G). By visual inspection, the addition of extra labels removed the largest and clearest misclassifications (Fig 7 F, G) and the ones remaining were overlapping with areas where the volume of training data was already large. As overall image detection parameters were not better for **RF-BF-2** compared to **RF-BF-1** (Table 1), it was deemed that the model quality had reached a plateau and further addition of data would not provide any significant model generalization.

### Cell segmentation with deep learning

For training the models for the DL pipeline, ten in-focus RFP fluorescence channel images of CHO-K1-hM_4_R cells stained with DiI dye were manually labeled using the ilastik Pixel classification pipeline user interface. For that, pixels were classified as either cells or background. The manually generated annotations were exported. Next, a background correction step was used to remove systematic illumination differences from the fluorescence images. The ten images along with corresponding ground truth annotations were randomly sampled into training, validation and test sets as follows: six images in the training set, two images in the validation set and two images in the test set. The training and validation set images were cropped to the input size of the U-net (288 x 288 pixels) and augmented using a sequential augmenter with the augmentations (rescaling 0 - 5%, shearing 0 - 1 pixels, piecewise affine shearing 1-5%, random rotation +-45 degrees, random left-right flip 50% probability and random up-down flip 50% probability) using the imgaug library [54]. A total of 6000 training tiles and 2000 validation tiles were generated (1000 augmented tiles of each image). The U-Net inspired fully convolutional U-Net3 architecture (Fig. 1 C) was used to train a model (**U-Net3-FL-1**) for cell detection from the fluorescence images [47, 49]. The training was carried out using the following parameters: Adam optimizer [55], learning rate=0.0002, beta 1=0.9, beta 2=0.999, epsilon=10^-8^, number of epochs=20, loss function=binary cross-entropy. The validation set loss was confirmed to have reached a minimum within 20 epochs. The model quality was assessed for the test set images. Next, the model was used to predict the masks for 191 DiI labeled fluorescence images. These images were again separated into training, validation and test sets along with the corresponding in-focus bright-field images of the same fields of view (133, 29 and 29 images respectively). The focal plane had been manually chosen in a prior step. As it has been previously shown that similar DL network architectures require considerably more bright-field data to converge to an optimal solution compared to fluorescence data, a different strategy was chosen for training DL for cell detection from bright-field images [40,49,56]. As the training data volume was substantially larger, a data generator was used for cropping the images to the correct size (288 x 288 pixels) instead of predefined training and validation sets. A batch size of 8 images was used during training. No augmentation was used for bright-field data. The same model architecture was used for the **U-Net3-BF-1** model as for **U-Net3-FL-1.** In this training run, early stopping with patience = 20 was used, the model converged after 90 epochs. Also, learning rate reduction with a factor of 0.1 and patience = 10 was used. All other parameters were the same as for the fluorescence-based model. The final model **U-Net3-BF-1** was used to predict the segmentation of the test set and equivalent metrics were calculated (Table 1).

**Fig. 1.**
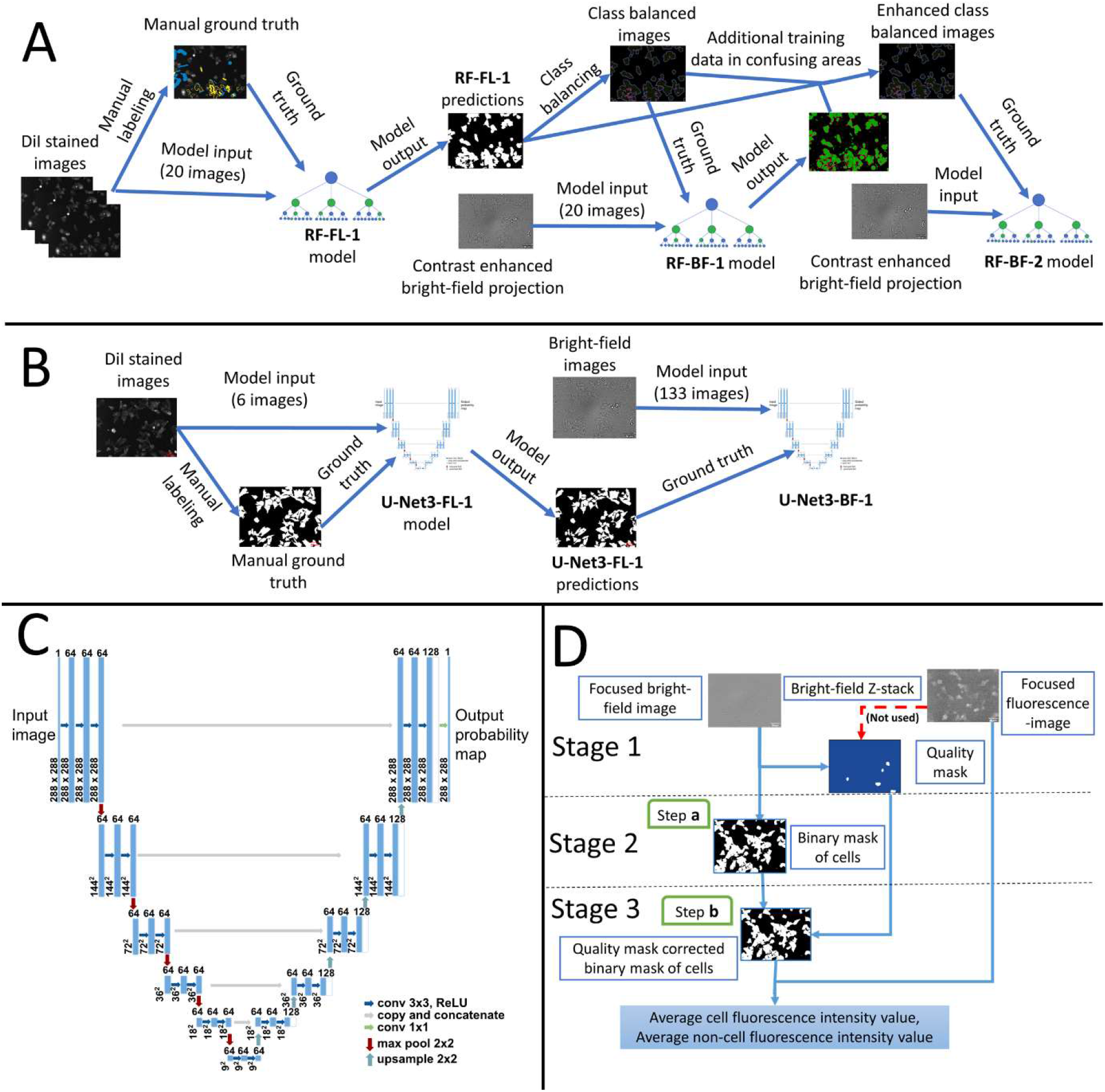
The architecture of the image analysis pipelines. For training the random-forest based **RF-BF-2** model, a training pipeline (A) was used which also generated the helper models **RF-FL-1** and **RF-BF-1**. For training the **U-Net3-BF-1** model the training pipeline (B) was used which also generated the helper model **U-Net3-FL-1.** U-Net type architecture (C) was used for the DL pipeline. The cell detection pipeline using the trained models (D) was implemented in Aparecium *MembraneTools* module. In this pipeline, **Stage 1** corresponds to data pre-processing, **Stage 2** to cell segmentation and **Stage 3** to data post-processing. In **Step a** of the cell detection pipeline, the **U-Net3-BF-1** model is used and in **Step b** the binary mask is corrected with the quality mask.

**Fig. 2.**
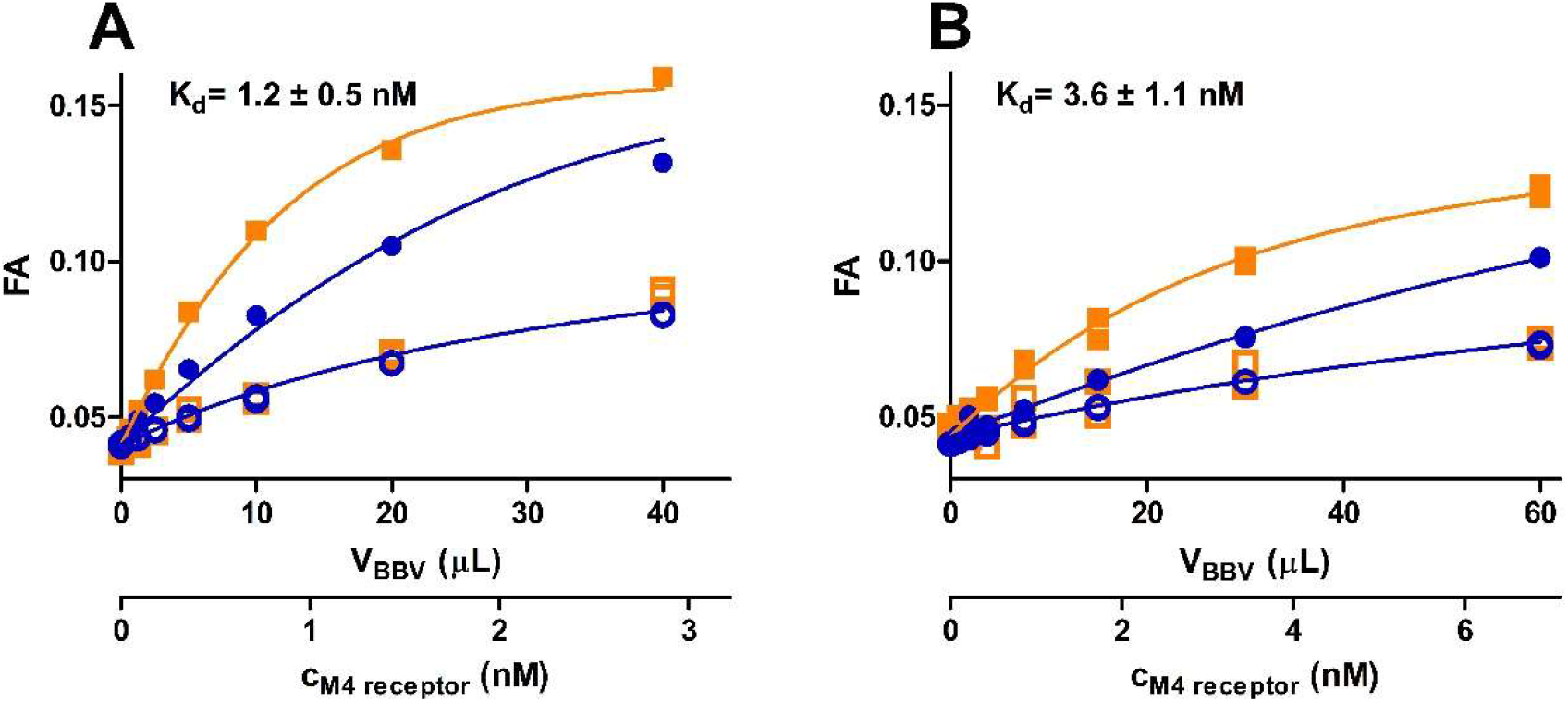
Binding curves of fluorescent ligand binding to M_4_ receptor. 1 nM 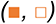 or 6 nM 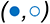 UR-MK342 (A) or 2 nM 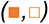 or 20 nM 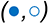 UR-CG072 (B) incubated with different concentrations M_4_ receptor. Total binding 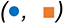 was determined in the absence and nonspecific binding 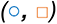 in the presence of 1000-fold excess of UNSW-MK259 in the case of UR-CG072 or scopolamine in the case of UR-MK342. The 1 h measurement point for UR-CG072 and 5 h measurement point for UR-MK342 are shown and was used for calculations. The concentration of M_4_ receptor binding sites (C_M4_) was calculated post hoc from the results of these experiments using the model described in [28]. A representative experiment of at least three independent experiments is shown. Experiments were performed in duplicates with all measurement data points shown.

### Image analysis pipeline

To carry out cell segmentation from all microscopy images, a suitable image analysis pipeline was developed. For using ilastik based models, the same pipeline was used as in [35] with minor modifications. The ilastik segmentation label index was updated according to the **RF-BF-1** model design (segmentation label index = 2) and morphological corrections were not utilized as it was not necessary for whole-cell segmentation in contrast to contour segmentation.

For using the **U-Net3-BF-1** model for prediction, the *MembraneTools* module of Aparecium data and image analysis software (https://gpcr.ut.ee/aparecium.html) was updated to be able to use Keras framework [57] models for prediction. Unlike for **RF-BF-1**, for **U-Net3-BF-1** only a single in-focus bright-field image was used for input instead of the contrast-enhanced image generated from bright-field Z-stacks. As the **U-Net3-BF-1** model can predict only the 288 x 288-pixel patches, the bright-field images are tiled before prediction and the predictions are later stitched to original size images (Fig. 1 D, step a).

The quality mask was manually generated (Fig 1 D, Stage 1) as previously described [35] and areas of low quality were removed from image quantification (Fig 1 D, step b).

For fluorescence image quantification, the in-focus fluorescence image was selected from the Z-stack manually and the image intensity was calculated only for the areas detected as cells by the segmentation model.

### Software performance tests

During software performance tests ilastik version 1.3.3post3 and MATLAB (The MathWorks, USA) version R2021a were used on a PC with 16 GB RAM, i7-10750H CPU (2.6 GHz) with 6 cores. DL pipeline inference was run on Nvidia Quadro T1000 GPU. For training the DL models the High Performance Cluster of University of Tartu was used [58].

### Pharmacological data analysis

Aparecium 2.0 software was used to blank the raw parallel and perpendicular intensity values and calculate the FA values using the formula [59]:

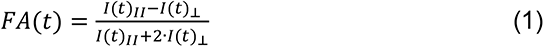

where, I(t)_II_ is the parallel fluorescence intensity, and I(t)_⟂_ is the perpendicular fluorescence intensity at time point t.

K_i_ values were calculated with the Cheng-Prusoff equation [60], using the IC_50_ values gained from data fitting with GraphPad Prism 5.0 (GraphPad Software, San Diego, USA) with a three-parameter logistic regression model (“log(inhibitor) vs. response”). For calculation of kinetic parameters k_on_, k_off_ and K_d_kinetic_ of the microscopy data, GraphPad Prism 5.0 “Association then dissociation” model was used. K_d_ calculation form microscopy data was also done with GraphPad Prism 5.0, but the model used was “One site --Total and nonspecific binding”.

For K_d_ calculation from FA data a global model form [28], which takes ligand depletion into account was used. To calculate k_on_, k_off_ and K_d_kinetic_ from FA kinetic data a modified version of IQMTools/SBToolbox2 (IntiQuan, Basel, Switzerland) was used to fit FA values with the previously published model [17]. The model assumes four possible interactions: the interaction between the receptor (R) and the fluorescence ligand (L), receptor and the competitive unlabeled ligand (C), nonspecific binding sites from the receptor preparation (NBV) and fluorescent ligand, the interaction between nonspecific binding sites on the microplate (N) and the fluorescent ligand. The corresponding reactions can be described by the following schemes:

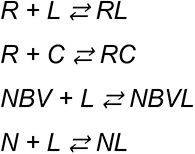

The concentrations in this model are connected to the predicted FA values through the equation:

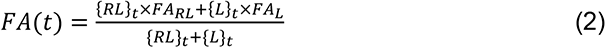

where *{RL}_t_* and *{L}_t_* are the instantaneous concentrations of *RL* and *L* respectively at timepoint *t* and *FA_RL_* and *FA_L_* are the intrinsic fluorescence anisotropies of the *RL* and *L* states respectively.

All the uncertainties given are weighted standard error of the mean of at least 3 independent experiments if not stated otherwise.

### Statistical analysis

For determining the quality of all machine learning cell detection models, four metrics were considered:

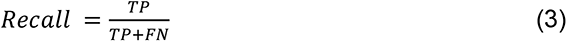

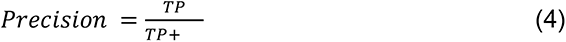

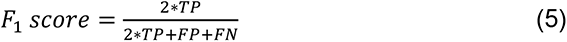

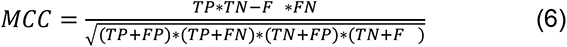

where True positive (TP) denotes the number of correctly detected pixels belonging to cells, True negative (TN) is the number of correctly predicted pixels not belonging to cells, False positive (FP) is the number of non-cell pixels detected as cells, False negative (FN) is the number of cell pixels detected as non-cell pixels.

To compare **U-Net3-BF-1** and **RF-BF-2** model qualities for determining IC_50_ values from the live-cell microscopy assay, the R^2^ of the nonlinear fits were compared in a pairwise manner using one-tailed Mann-Whitney U-test in GraphPad Prism 5.0 assuming that **U-Net3-BF-1** is the superior model.

To determine the assay suitability for HTS applications, Z’ values were calculated according to the formula [61]:

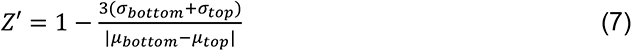

Where *σ_top_* and *σ_bottom_* represent the standard deviations of blank wells (negative controls) and wells with a full displacement of the fluorescent ligand (positive controls), respectively. *µ_top_* and *µ_bottom_* correspond to means of negative and positive controls, respectively. The values from individual experiments were normalized to the respective top and bottom plateau values for each concentration-response curve separately to remove batch-to-batch variation effects of the receptor source.

## 3. Results

### Determination of binding affinities of UR-CG072 and UR-MK342 to M4 receptor displayed in BBV particles with FA

The fluorescence anisotropy method used here allows continuous measurement of receptor-fluorescence ligand complex formation or dissociation. FA depends on the ratios of free and bound fluorescence ligand states (equation 2). The specific fluorescence anisotropy values of each state depend on the fluorescence lifetime as well as the rotational freedom of the fluorophore in the corresponding state. To observe significant changes in FA signal it is necessary that the mole ratios of both the free and bound fluorescence ligand change when receptor concentration, total fluorescence ligand concentration, competitive ligand concentration, time or a combination of these factors is varied. This is best achieved when concentrations of the probe and its target protein are kept close to their binding K_d_.

For these reasons only certain fluorescence ligands with suitable fluorophores and binding affinities, usually from low picomolar to low nanomolar ranges, are considered as probe candidates for FA assay. Two 5-TAMRA labeled ligands, UR-CG072 and UR-MK342 from [15], were chosen for the development of fluorescence anisotropy assays due to a suitable label and high-affinity to M_4_ receptor determined by radioligand binding to whole cells.

First, the saturation binding experiments were carried out to determine fluorescence ligand binding affinities to M_4_ receptors displayed on BBV particles. Both ligands showed similar and high binding affinity (K_d_UR-CG072_ = 3.6 ± 1.1 nM, K_d_UR-MK342_ = 1.2 ± 0.5 nM) which are in good agreement with the radioligand binding values (K_i_UR-CG072_ = 3.7 ± 0.6 nM, K_i_UR-MK342_= 0.97 ± 0.07 nM [15]). However, UR-MK342 binding has a larger dynamic range of FA values compared to UR-CG072. The same tendency was also found in FA assays with the M_2_ receptor [17]. As the effect is evident for both receptor subtypes, it might be attributed to the more flexible linker in UR-CG072. Nevertheless, high-affinity and sufficient dynamic range mean that both ligands would be suitable for kinetic measurements as well as using these as probes for measuring competitive ligand binding parameters.

First, the saturation binding experiments were carried out to determine fluorescence ligand binding affinities to M_4_ receptors displayed on BBV particles. Both ligands showed similar and high binding affinity (K_d_UR-CG072_ = 3.6 ± 1.1 nM, K_d_UR-MK342_ = 1.2 ± 0.5 nM) which are in good agreement with the radioligand binding values (K_i_UR-CG072_ = 3.7 ± 0.6 nM, K_i_UR-MK342_ = 0.97 ± 0.07 nM [15]). However, UR-MK342 binding has a larger dynamic range of FA values compared to UR-CG072. The same tendency was also found in FA assays with the M_2_ receptor [17]. As the effect is evident for both receptor subtypes, it might be attributed to the more flexible linker in UR-CG072. Nevertheless, high-affinity and sufficient dynamic range mean that both ligands would be suitable for kinetic measurements as well as using these as probes for measuring competitive ligand binding parameters.

Next, the ligand binding kinetics of UR-CG072 and UR-MK342 to the M_4_ receptor were studied. In contrast to similar affinities, the kinetic properties of UR-CG072 and UR-MK342 were quite different (Fig. 3, Table 2). The faster association and dissociation kinetics of UR-CG072 make it more suitable for FA based screening assays as this allows increasing the assay throughput by shortening the incubation times which reduces problems concerning potential receptor source sedimentation, liquid evaporation or even degradation of the ligands or the receptor [62]. Faster kinetics is also beneficial for live-cell microscopy assays, where too long experiments can lead to problems with cell culture such as detachment and changes in medium composition.

**Fig. 3.**
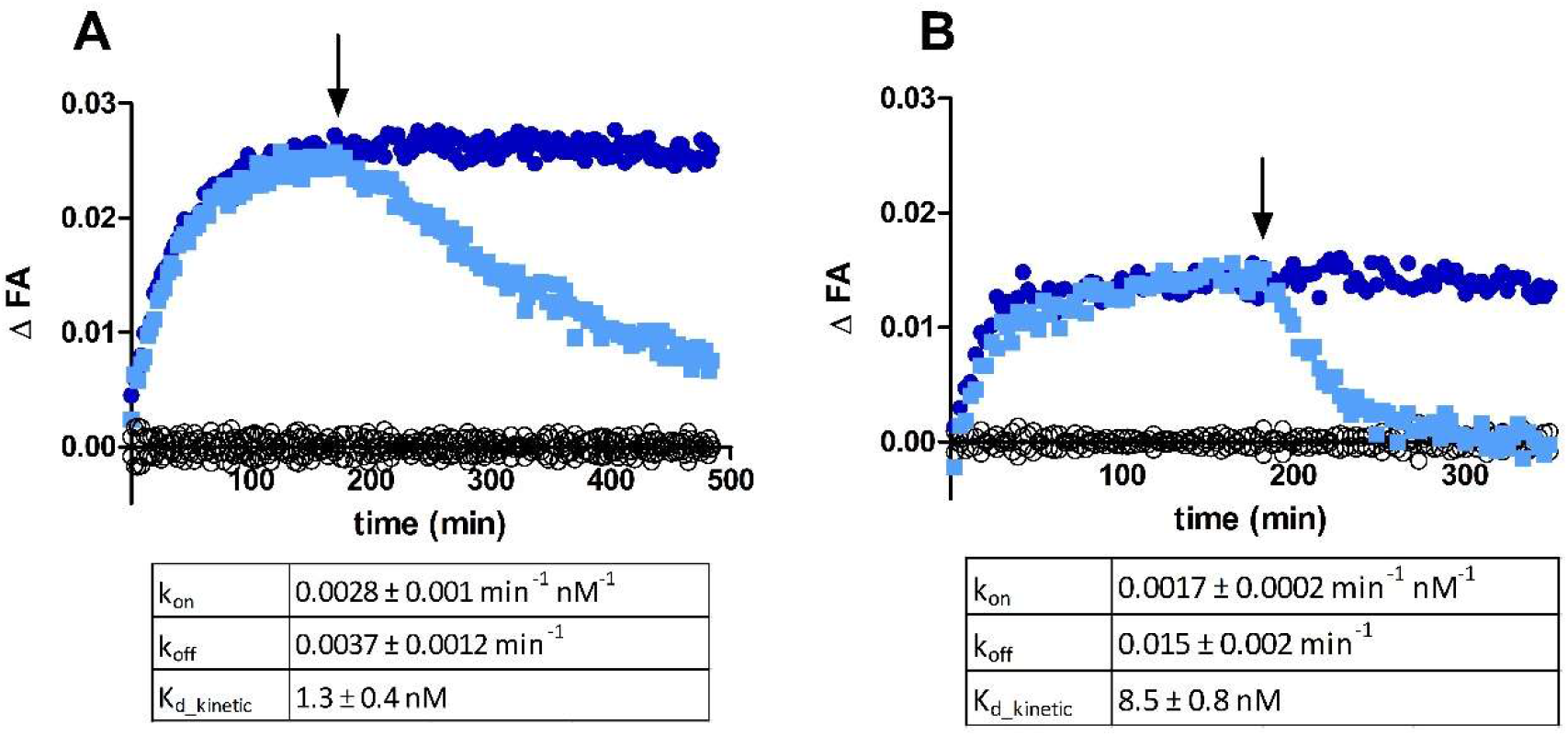
Time course of FA change caused by UR-MK342 (A) or UR-CG072 (B) binding to M_4_ receptors on the BBV particles. The reaction was initiated by the addition of 20 μL M_4_ receptor displaying BBV particles to 6 nM UR-MK342 (A) or 5 nM UR-CG072 (B) in the absence 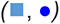 or presence (**○**) of 6 μM (A) or 3 μM (B) scopolamine, respectively. After 180 min (indicated with an arrow) dissociation was initiated by the addition of 6 μM (A) or 3 μM (B) scopolamine 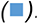. An equivalent volume of assay buffer was added to association controls 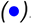. Representative experiments of at least three independent experiments are shown. ΔFA is calculated by subtracting the FA value of nonspecific binding from the measured FA value of the corresponding measurement.

**Table 2.**
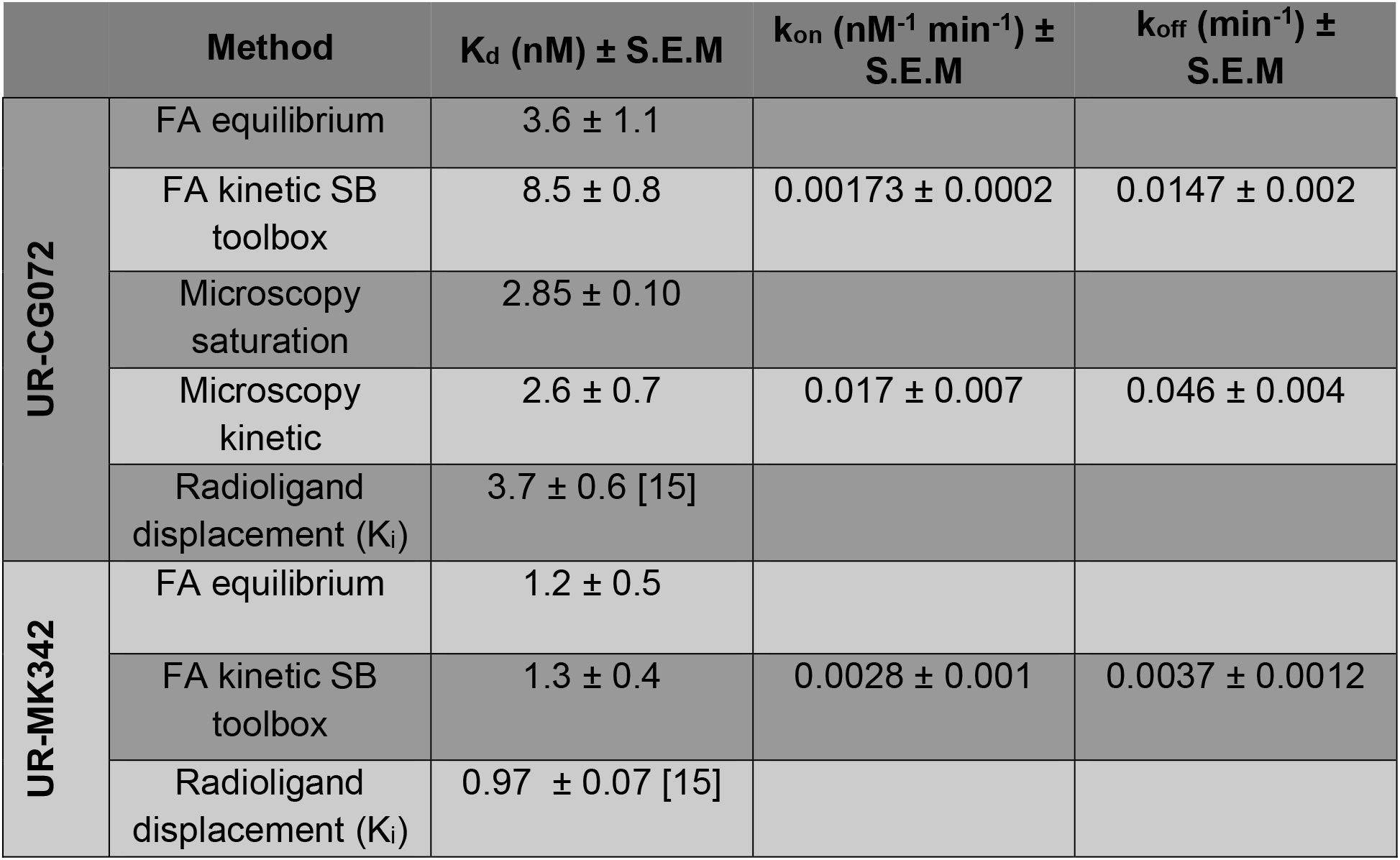
Overview of binding parameters of UR-CG072 and UR-MK342

| | Method | $K_d$ (nM) ± S.E.M | $k_{on}$ (nM<sup>-1</sup> min<sup>-1</sup>) ± S.E.M | $k_{off}$ (min<sup>-1</sup>) ± S.E.M |
| --- | --- | --- | --- | --- |
| UR-CG072 | FA equilibrium | $3.6 \pm 1.1$ | | |
| | FA kinetic SB toolbox | $8.5 \pm 0.8$ | $0.00173 \pm 0.0002$ | $0.0147 \pm 0.002$ |
| | Microscopy saturation | $2.85 \pm 0.10$ | | |
| | Microscopy kinetic | $2.6 \pm 0.7$ | $0.017 \pm 0.007$ | $0.046 \pm 0.004$ |
| | Radioligand displacement ($K_i$) | $3.7 \pm 0.6$ [15] | | |
| UR-MK342 | FA equilibrium | $1.2 \pm 0.5$ | | |
| | FA kinetic SB toolbox | $1.3 \pm 0.4$ | $0.0028 \pm 0.001$ | $0.0037 \pm 0.0012$ |
| | Radioligand displacement ($K_i$) | $0.97 \pm 0.07$ [15] | | |

### Affinity screening a panel of MR ligands with UR-CG072 and UR-MK342

Fluorescence ligands are often applied for determining the affinities of unlabeled ligands. Therefore, the suitability of both ligands was studied as reporter probes in competition with unlabeled M_4_ receptor ligands. For that, a panel of common M_4_ receptor ligands was chosen such that the expected affinities would cover a wide range of values and contain both agonists and antagonists. In addition, some unlabeled ligands, which are structurally similar to the fluorescence ligands [50,51,63], were chosen to assess the assay’s ability to work with dualsteric compounds. The set of ligands was investigated in competition binding experiments with both UR-CG072 and UR-MK342 (Fig. 4).

**Fig 4.**
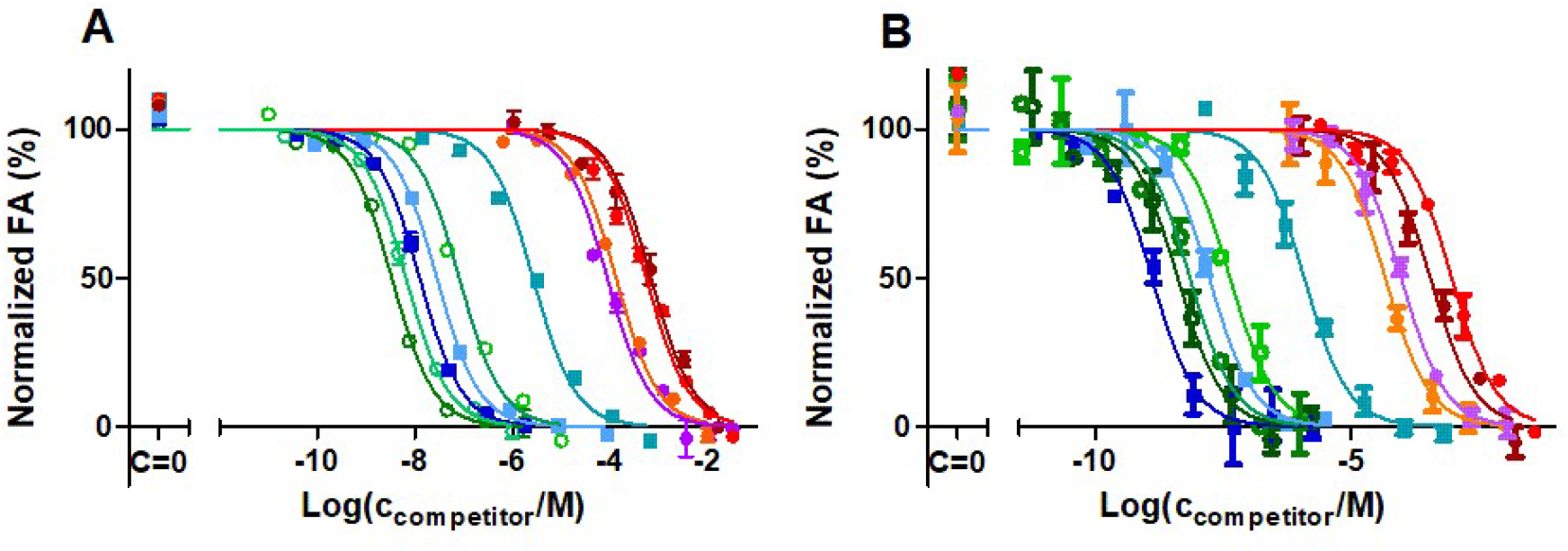
FA-based M_4_ receptor competition binding experiments performed with 5 nM UR-MK342 (A) or 5 nM UR-CG072 (B) and reported muscarinic M_4_ receptor ligands. BBV particles displaying the M_4_ receptors (V(BBV) = 20 μL) were used as the receptor source. The 9 h measurement point is shown and used for analysis for all ligands. From the used competitive ligands, acetylcholine 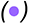, carbachol 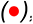, arecoline 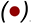, pilocarpine 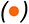 are agonists, scopolamine 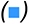, atropine 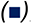, pirenzepine 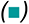, UNSW-MK259 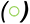, UR-SK75 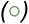 and UR-SK59 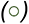 are antagonists. A representative experiment of at least three independent experiments performed in duplicates is shown. The error bars represent the SEM of duplicates. Normalization was performed by taking the upper plateau value as 100% and lower plateau value as 0% separately for each displacement curve.

Both fluorescent ligands can successfully be used as reporter ligands with a high signal-to-noise ratio and very good Z-prime (Z’_UR-CG072_ = 0.52, Z’_UR-MK342_ = 0.67) making the assay compatible with high-throughput screening formats which generally require a minimum Z’ of 0.5. However, as UR-CG072 has faster kinetics than UR-MK342, a longer incubation time is needed to determine the competitive ligand affinity using UR-MK342. To avoid possible under-or overestimation of IC_50_ values, it is important to wait until the equilibrium is reached [22].

To make the measurement values comparable, the pK_i_ values were calculated from the IC_50_ value for each ligand using the Cheng-Prussoff equation. While not all assumptions of the Cheng-Prussoff equation [60] are fulfilled, it has been previously shown that with these ligands the potential systematic error introduced by this operation is relatively small [17]. pK_i_ values obtained from experiments using the two different reporter ligands correlated very well (R^2^ = 0.96) and the linear regression slope of the obtained pKi values with both probes is very close to unity (0.97 ± 0.04) while the intercept is close to zero (0.3 ± 0.3) (Fig. 5 A). This validates that both probes can be used to determine the unlabeled ligand affinities in the FA assay.

**Fig 5.**
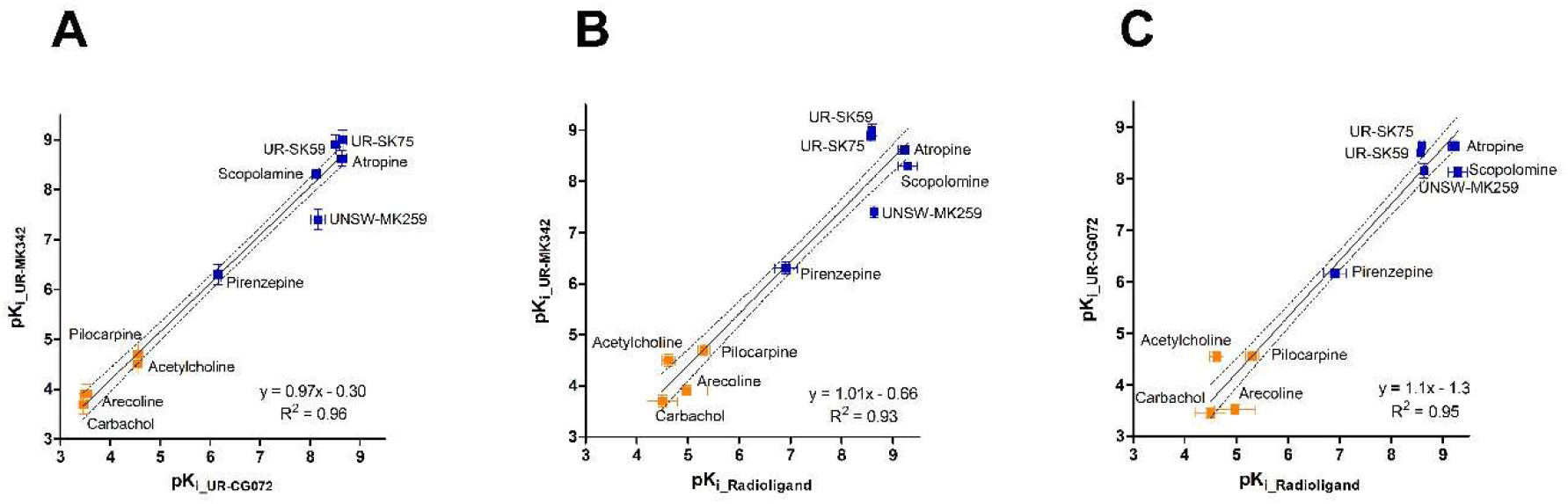
Correlations of binding affinities (pK_i_ values) of ligands to M_4_ receptor, measured with different probes and assays. (A) FA assays with UR-CG072 and UR-MK342; (B) FA assay with UR-MK342 and radioligand binding (literature data); (C) FA assay with UR-CG072 and radioligand binding (literature data); Investigated agonists are presented as orange symbols 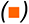, antagonists as blue symbols 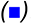. Black lines represent the best linear relationship between the datasets and the dashed black lines represent the 95% confidence bands.

Out of the tested ligands, UNSW-MK259, which represents non-labeled analogue of UR-CG072 (Fig 5), had the largest deviation from the best regression line. The reason for this deviation is unknown but may be connected to potential dualsteric binding modes of UNSW-MK259, UR-MK342 and UR-CG072 which could alter the binding mechanism. However, explaining this effect remains the topic of future studies.

### Adjusting live cell microscopy assay for measuring UR-CG072 binding to M4 receptor

To keep the cells viable and with normal morphology during imaging experiments it is necessary to maintain specific conditions, like 5% CO_2_, 37 °C and sufficient nutrient concentrations in the media. These parameters may start to drift over long periods. Therefore, UR-CG072 was selected for the live-cell assay, due to its faster binding and dissociation kinetics. First, it was confirmed that the binding of 2 nM UR-CG072 to CHO-K1-hM_4_R cells can be detected by fluorescence microscopy (Fig. 6 A1). For nonspecific binding controls, a similar experiment was performed in the presence of 5 µM scopolamine. As illustrated in Fig 6 there is a significant difference in fluorescence intensity between total binding (Fig. 6 A1) and nonspecific binding (Fig. 6 B1). To confirm that all the signal is specifically caused by ligand binding to M_4_ receptors, the binding of 2 nM UR-CG072 to CHO-K1 cells not expressing M_4_ receptor was measured. Under these conditions, there was no detectable accumulation of UR-CG072 to CHO-K1 cells (Fig. 6 C1).

**Fig. 6.**
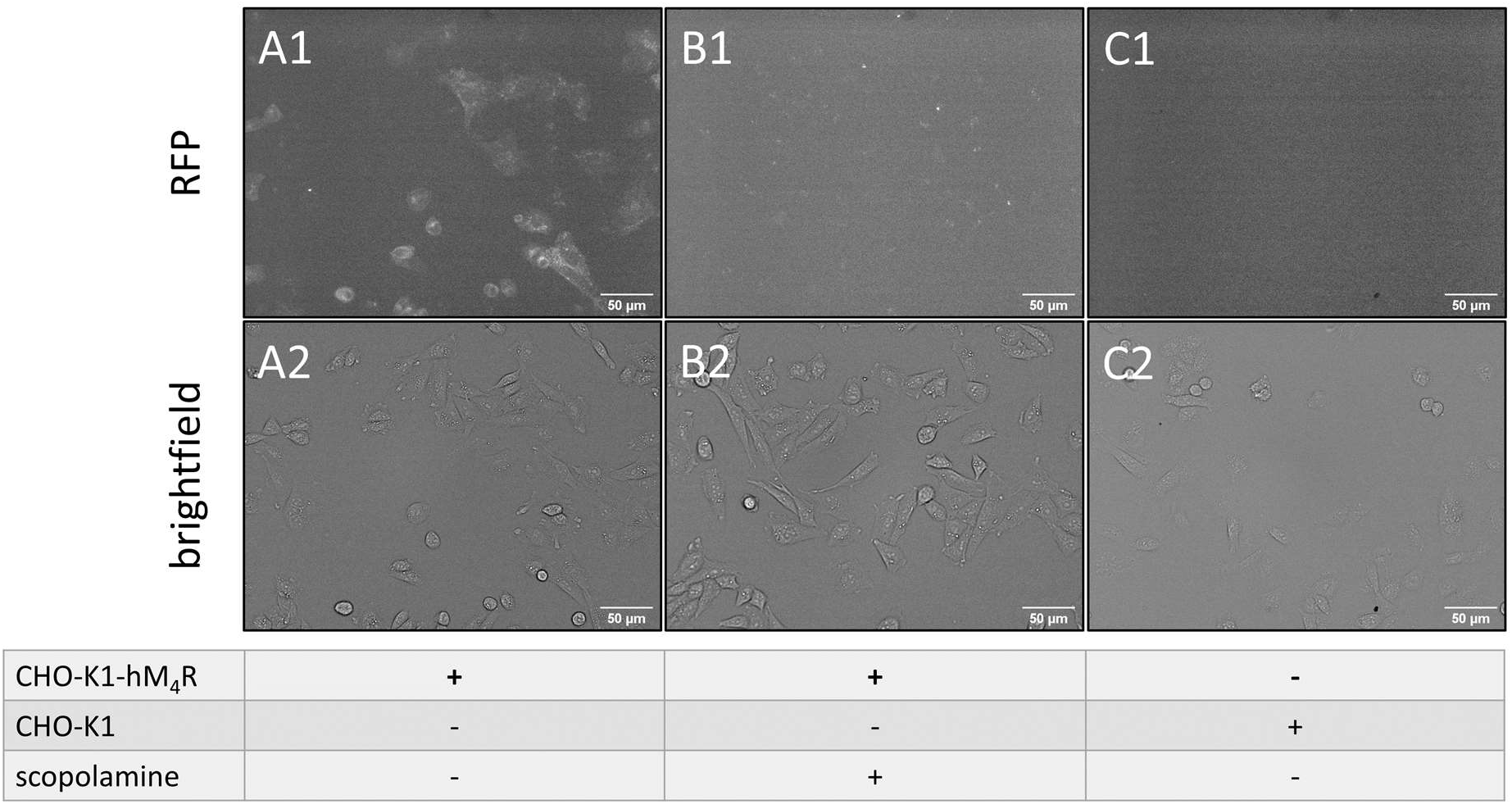
Fluorescence microscopy images (top row) and corresponding bright-field images (bottom row) of UR-CG072 binding to CHO-K1 cells. Either wild-type CHO-K1 cells (C1 and C2) or CHO-K1-hM_4_R cells (A1, A2, B1, B2) in DMEM/F12 medium with added 9% FBS and antibiotic antimycotic solution were incubated with 2 nM UR-CG072 in the absence (A1, A2, C1, C2) or presence (B1, B2) of 5 μM scopolamine for 3 h. The number of seeded cells per well was 30 000. The contrast of fluorescence images was enhanced for presentation purposes only, the same lookup table was used for all images. The scale bar corresponds to 50 μm.

**Fig. 7.**
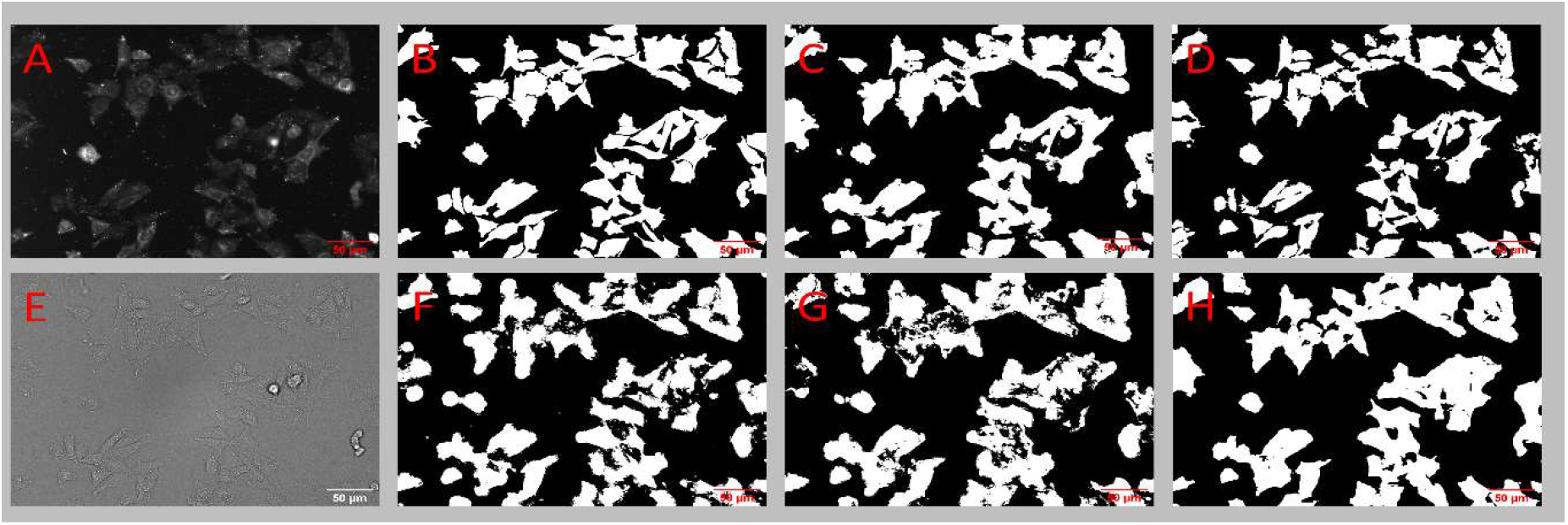
Microscopy images and corresponding binary masks of CHO-K1-hM_4_R cells stained with DiI. The top row shows cell detection from the fluorescence image (A) by manual segmentation (B), **RF-FL-1 model** (C) and **U-Net3-FL-1** (D). The bottom row shows cell detection from the bright-field image (E) by **RF-BF-1** model (F), **RF-BF-2** model (G) and **U-Net3-BF-1** (H). The scale bar corresponds to 50 µm.

The results show that the differences between cell contour and cell body fluorescence intensities are smaller for the flatter CHO-K1 cells compared to HEK293 cells used in a previous study [35], which are elongated in the Z-direction. Therefore, it is necessary to analyze the fluorescence intensity of the whole cell body which introduces an increased proportion of cell autofluorescence to the signal. Moreover, the imaging experiments were carried out in nutrient-rich cell culture media rather than DPBS buffer as is suggested in [35]. This removes the need for more expensive special imaging media but increases background fluorescence levels. Combining these effects with using 5-TAMRA fluorophore instead of Cy3B as was used in [35], the overall signal level was greatly reduced in this assay compared to the previous one. However, as biological variability is still the main contributor to the assay uncertainty, reducing such variability at the cost of a reduced signal is still beneficial to the overall assay quality.

These images also reveal that the amount of M_4_ receptors on the cell membrane surface in all cells is different and that in some cells ligand binding could not be detected at all (Fig. 6, A1-A2). This aspect should be considered when moving on to single-cell based quantification using this particular cell line. However, the current microscopy method averages the signal from a large number of cells and all cells used on a single assay day are seeded from the same population. Furthermore, the absolute intensity values have no direct or systematic influence on the calculated ligand binding parameters (K_d_, k_on_, k_off_ and K_i_) and only lead to lowered signal-to-noise ratio. While a higher signal-to-noise ratio is beneficial in general, in the current case, it only has a limited impact on the overall measurement uncertainty.

### Comparison of Random Forest and Deep learning-based image analysis pipelines

Since the morphologies of HEK293 cells and CHO-K1 cells and the fluorescence probes are different, it was necessary to adjust the original pipeline previously developed for HEK293 cell analysis [35]. Due to a lower contour contrast of CHO-K1 cells compared to HEK293 cells, it was necessary to quantify the fluorescence intensity from the entire cell mask instead of only the cell contours.

A second adjustment to the original pipeline was needed due to the lower apparent brightness of the ligand-receptor complex. While the NAPS-Cy3B fluorescence signal in the dopamine D_3_ receptor system was close to twice as high as the image background intensity [35], the signal of CHO-K1-hM_4_R bound UR-CG072 was only 4% above the background signal. Due to the high absolute signal level in the D_3_ receptor system, it was not necessary to find the in-focus fluorescence plane in the original pipeline and instead the maximum intensity projection of the Z-stack could be used. In the current case, this approach is not suitable and leads to complete signal degradation (data not shown). Therefore, the fluorescence intensity must be quantified from the highest quality focal plane.

Improvements were also introduced into the model development and ground-truth generation process. The original pipeline relied on a human analyst to detect cell contours from bright-field images. Even though it was necessary to perform this step only once, it required significant manual labour and detecting cell contours from bright-field images is still more difficult compared to detection from fluorescence images. To address these issues, another approach was pursued. The cell membranes were stained with a lipophilic dye DiI and then imaged in both fluorescence and bright-field channels. For a small number of fluorescence images, cell masks were manually drawn. Next, machine learning models **RF-FL-1** and **U-Net3-FL-1** were trained to generate cell masks from the DiI stained fluorescence images. These models were in turn used to predict the masks from a larger dataset of fluorescence images. The prediction masks then served as slightly lower quality, but significantly higher quantity ground truth for the next set of models (**RF-BF-1**, **RF-BF-2** and **U-Net3-BF-1**) which predict the cell masks from bright-field images. The same conceptual approach was successful for training both the RF-based pipeline implemented in ilastik as well as the U-Net3 based deep-learning pipeline developed using Jupyter notebooks [64] and Keras deep-learning framework [57]. Considering all the aspects, both developed pipelines were superior to the original pipeline from the pipeline development perspective with a significantly reduced amount of manual annotation required.

### Prediction quality comparison

The prediction quality of the deep-learning models and ilastik based RF model were compared to determine the most suitable pipeline for analysis (Fig. 7, Table 1). Visually, all models can segment most of the cells from the bright-field images with good quality. The main difference between the models is that **U-Net3-BF-1** (Fig. 7 H) produces cells with more consistent and smooth shapes, similar to the ground truth (Fig. 7 B) while **RF-BF-2** (Fig. 7 G) creates rugged edges and also detects many small fragmented objects far from the cells. Numerically, the quality of bright-field detection (Table 1, F1 score = 0.89) is somewhat lower than the current state of the art solutions in the cell tracking challenge [65] when compared to the most similar dataset of DIC-HeLa cells (F1 score = 0.93) [65–68]. However, it must be noted that such small differences could be easily caused by differences in imaging modes, magnifications, cell line morphology, amount of training data or among other parameters. The F1 score of the fluorescence image-based predictions of both **U-Net3-FL-1** (Fig. 7 D) and **RF-FL-1** (Fig. 7 C) are already substantially lower than unity. At the same time, the **U-Net3-BF-1** model has only slightly lower quality metrics compared to **U-Net3-FL-1** but **RF-BF-2** has substantially lower metrics compared to **RF-FL-1**. This may indicate that a large proportion of the errors made by DL pipeline originates from the training of the fluorescence model **U-Net3-FL-1** rather than the bright-field model **U-Net3-BF-1**. Interestingly, when comparing the **U-Net3-FL-1** model predictions and the **U-Net3-BF-1** model predictions directly to one another instead of comparing these to the manually generated ground truth, the corresponding F1 score is 0.87. This is lower than the similarity between either the **U-Net3-FL-1** model and manual ground truth (F1 score = 0.91) or **U-Net3-BF-1** and manual ground truth (F1 score = 0.89). It means that the **U-Net3-BF-1** model can surpass the prediction quality of the **U-Net3-FL-1** model predictions in some instances while failing to do so in other cases. The ability of **U-Net3-BF-1** to avoid at least some of the mispredictions generated by the **U-Net3-FL-1** model could mean that the proposed strategy of bright-field model generation is likely to work even with relatively small manually annotated datasets without the risk of overfitting. Interestingly, the **RF-FL-1** model has a higher F1 score and MCC value compared to **U-Net3-FL-1** model. However, these numbers should not be used to make conclusions about the general power of a particular machine-learning approach, since the training sets for models were not identical. Different training sets were used for practical considerations. For example, the datasets were chosen to be small enough that would allow training the models within a few hours and without the need for unconventionally large computational resources, while still achieving sufficiently high quality.

Furthermore, analyzing the competition, saturation and kinetic experiments, with both **U-Net3-BF-1** and **RF-BF-2** models provides the opportunity to compare the pipeline performances not only on the image level but also on the pharmacological level. As the most commonly used metric for fit quality the R^2^ values of the nonlinear model from each experiment were compared in a pairwise manner. The analysis revealed that the R^2^ values obtained from the DL pipeline are statistically significantly higher compared to the RF pipeline (p=0.03) calculated as described in Methods. **U-Net3-BF-1** based cell detection had a higher average R^2^ values (mean = 0.93±0.05 and median = 0.939) compared to the **RF-BF-2** based cell detection pipeline (mean = 0.89±0.09 and median = 0.911). The relatively large standard deviation of the R^2^ values shows that the algorithmic uncertainty is not the primary source of uncertainty and instead the variability is caused by biological factors. The high average R^2^ values indicate that both pipelines work well in general and the difference is not very large in absolute terms, but also that the small inaccuracies in the cell segmentation stage are not cancelled out during the post-processing steps. Instead, the errors are carried over and degrade the final fitting quality. Therefore, the U-Net3 based DL pipeline can still offer considerable advantages over RF based approach at both image level and downstream nonlinear regression level. Thus, from the quality perspective, it is reasonable to prefer the DL pipeline with **U-Net3-BF-1** over the RF pipeline using the **RF-BF-2** model. As the **U-Net3-BF-1** model showed higher overall quality, all the following presented results were obtained using the DL pipeline.

### Usability of Deep Learning and ilastik pipelines for microscopy image analysis

In addition to model quality, the usability aspects of the developed pipelines were compared. The most relevant ones were general computational hardware requirements, pipeline speed, convenience of using the pipelines in terms of user interfaces and finally, the convenience of developing new machine learning models in case of adapting the developed assay for a different microscope or cell line.

It was identified that the speed of the ilastik based RF models is substantially slower compared to the U-Net3 based DL models used for analysing the microscopy images. The difference was especially evident in the case when a GPU (graphical processing unit) was used for computations, which considerably speeded up the DL models. A modern computer was able to analyze the results with both DL and ilastik pipelines in a comparable time for preparing an experiment or performing the imaging, thus, making the analysis quite manageable. On average, analyzing a single 904 x 1224 pixel image took 12 seconds with RF pipeline and 3.5 seconds with DL pipeline.

Compared to spectroscopy methods, large data volumes generated by the microscopy experiments may cause storage issues. Therefore, before using the proposed microscopy methods, the user should make sure that sufficient memory is available for the experiments.

Another aspect to consider is the analysis convenience, which in the case of image analysis software is related to the need of manually adjusting the algorithm parameters and performing some of the image analysis, preprocessing or post-processing steps manually. For both DL and ilastik pipelines, no manual parameter adjustment is needed removing one common obstacle in image analysis. In addition to choosing convenient machine learning models, it was necessary to choose a suitable interface for using the machine learning models and performing the pre and post-processing steps. Many such interfacing software tools such as FIJI (DeepImageJ [69]), CellProfiler [70] and ImJoy [71] allow almost unlimited flexibility for developing image analysis pipelines but also require that users have some knowledge of how image analysis pipelines work internally. These software currently also do not provide convenient out-of-the-box options for metadata handling required for pharmacological assays. Therefore, we chose Aparecium software (https://gpcr.ut.ee/aparecium.html) as the interfacing platform, as it is specifically designed for making image analysis pipelines as user-friendly as possible through graphical user interfaces (GUI) while providing enough options for post-processing and metadata handling to carry out the biochemical analysis at the cost of less flexibility for general image analysis.

Finally, the aspect of machine learning model development was considered as it is usually necessary to retrain the models from scratch or perform transfer learning if the method is used for widely different datasets [72, 73]. In this study, two quite different model development environments were used. Model development in ilastik is relatively straightforward, requiring no programming skills and is done entirely through a GUI provided by the standalone ilastik software. Installing the software is very simple and there are multiple tutorials available for using the GUI. Development of the DL models including U-Net is somewhat more difficult, requiring access to a python installation and preferentially to a Jupyter notebook server. However, this process is significantly simplified thanks to the recently developed ZeroCostDL4Mic framework [72]. ZeroCostDL4Mic reduces the training process to a point-and-click level without the need to adjust the code. Therefore, both ilastik and deep-learning image analysis pipelines are sufficiently simplified that model training does not require extensive past experience with ilastik being the simplest option. Therefore, ilastik pipeline and the RF model is recommended for machine learning applications where ease-of-use is more important than a slight loss in quality. These practical considerations are quite dynamic as software tools develop and are likely to change in the future.

### Determination of binding affinities with UR-CG072 to M4 receptor in live cell microscopy

For determining the binding affinity of UR-CG072, an assay design similar to the radioligand saturation binding experiment was used. From these data a K_d_ of 2.85 ± 0.10 nM was obtained (Fig. 8), which is also in good agreement with all previous results (Table 2). Interestingly, there is a small decline in nonspecific binding with increasing concentration (Fig. 8), but it is not of biological origin and is instead explained by a shadow-imaging effect which is caused by nonspecific binding of UR-CG072 to the well surface making the background brighter than the cells. This effect, however, does not interfere with the overall measurement.

**Fig. 8.**
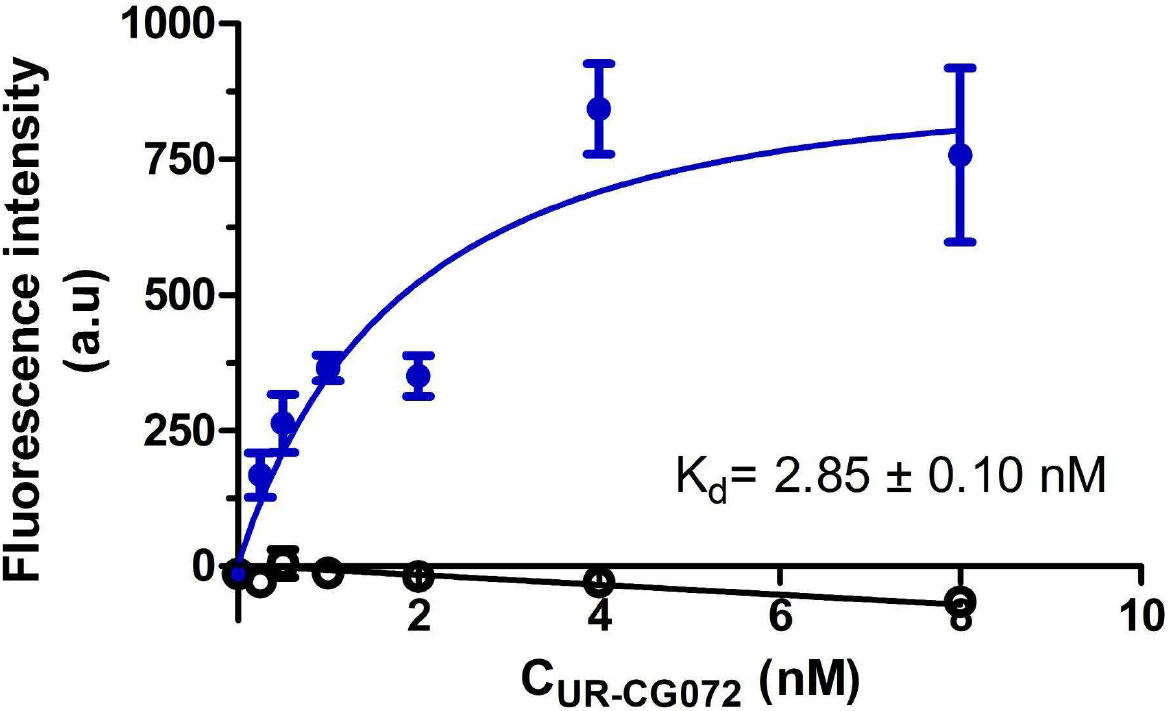
Saturation binding of UR-CG072 binding to live CHO-K1-hM_4_R cells. The CHO-K1-hM_4_R cells (25000 cells/well) were incubated for 5 h with two-fold serial dilutions of UR-CG072 in the range of 0–8 nM and with (nonspecific binding, 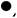) or without (total binding, 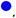) 3.7 µM scopolamine. The background-corrected cells’ fluorescence intensities were determined by the cell detection algorithm as described in Materials and Methods and are presented as mean ± SEM from a representative experiment of three independent experiments performed in duplicate. Four images from different fields of view were taken from a single well.

Due to the good photostability of the 5-TAMRA label and the moderate kinetic rates of UR-CG072 binding, the k_on_ and k_off_ of UR-CG072 could be measured with the described live-cell system. The binding of UR-CG072 (Fig. 9) is fully reversible by the addition of 10 µM scopolamine after 3 hours of association (indicated by the arrow). Moreover, the K_d_ (2.6 ± 0.7 nM) obtained from kinetic data is in good agreement with previous values from both saturation binding assay as well as fluorescence anisotropy assays (Table 2).

**Fig. 9.**
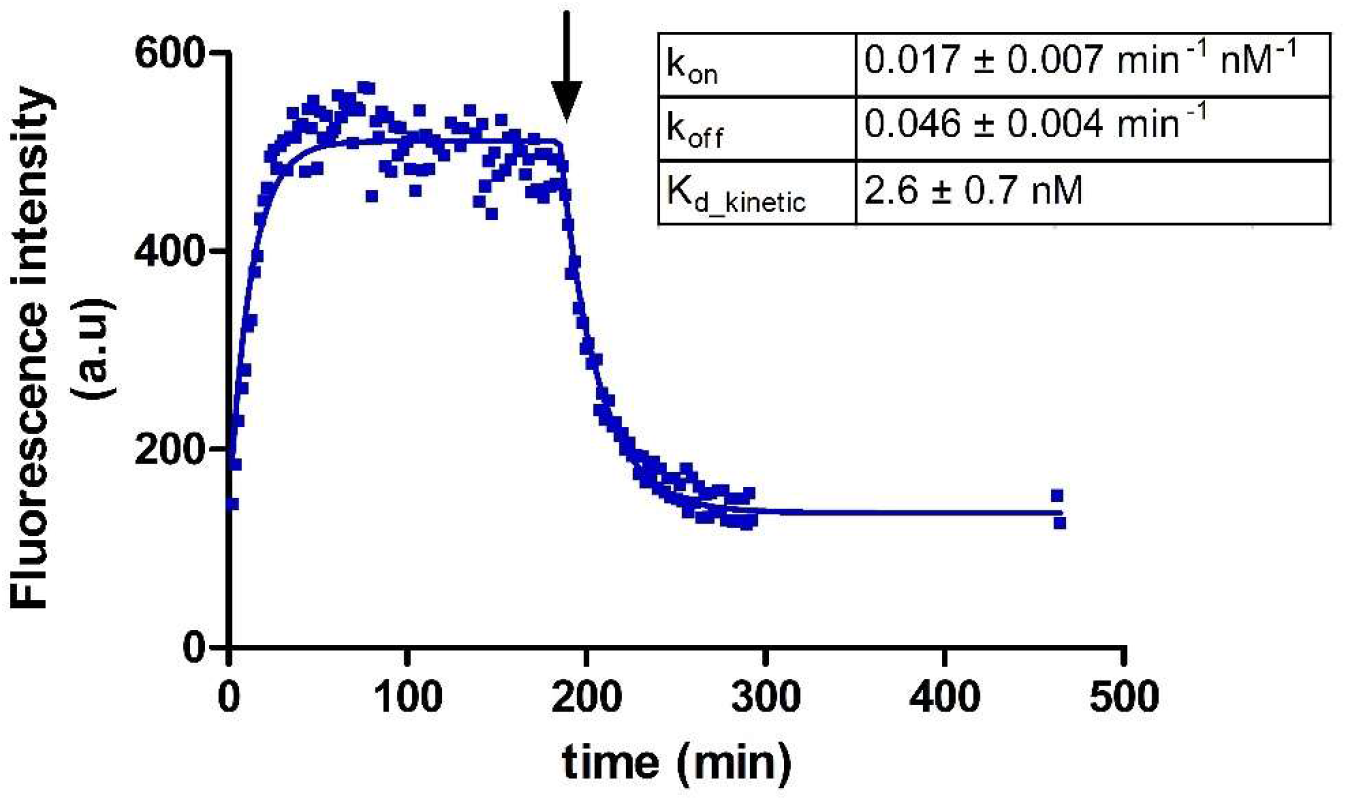
Association and dissociation of UR-CG072 (2 nM) to and from live CHO-K1-hM_4_R cells. Dissociation was initiated with 5 µM scopolamine after 185 min (indicated with the arrow). A representative experiment of three independent experiments performed in duplicates is shown. Duplicate values are shown on the graph as separate points, to account for the measurement time difference between the replicates. Four images from different fields of view were taken from a single well. For analysis the GraphPad Prism model “Association then dissociation” was used.

Lastly, competition binding experiments were carried out to confirm that the developed microscopy method is also suitable for screening novel unlabeled ligands in the future. Displacement curves were obtained for six ligands with varying structures, affinities and efficacies (Fig. 10). Regression analysis was used to obtain the IC_50_ values from these data, which in turn were used to calculate pK_i_ values of the unlabeled ligands (Table 3).

**Fig. 10.**
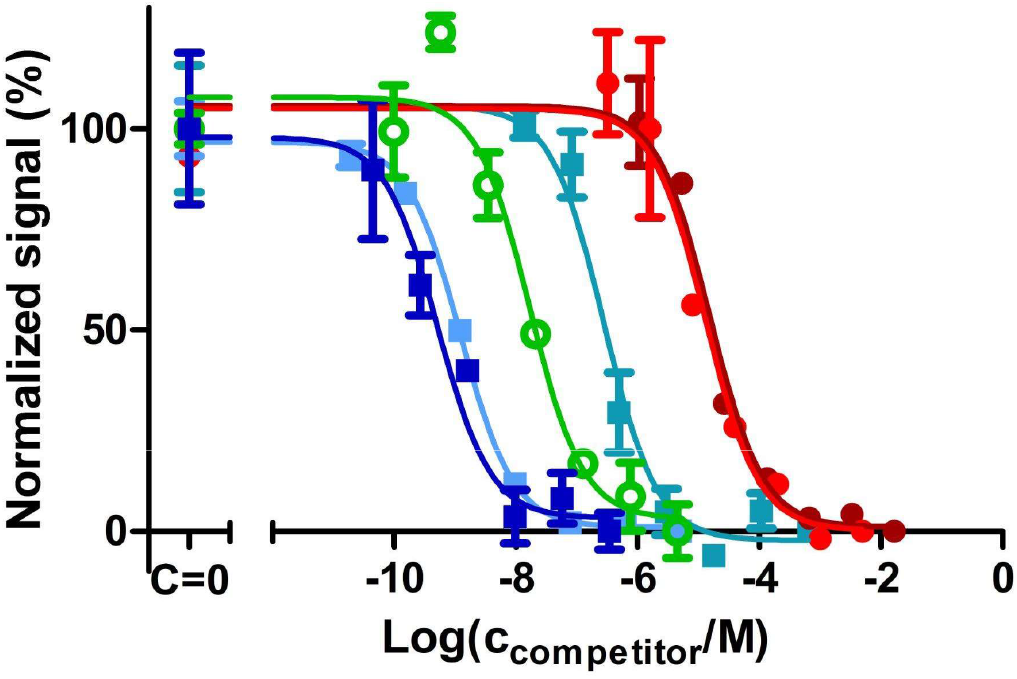
Inhibition of UR-CG072 binding to live CHO-K1-hM_4_R cells by muscarinic receptor ligands. The CHO-K1-hM_4_R cells (25 000 cells/well) were incubated with 2 nM UR-CG072 and different concentrations of carbachol 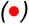, arecoline 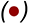, scopolamine 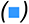, atropine 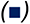, pirenzepine 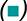, UNSW-MK259 for 120 min as described in Materials and Methods. The cell fluorescence intensities were determined by the membrane detection algorithm using the U-Net3-BF-1 model and are presented as mean ± SEM from a representative experiment performed in duplicate. Four images from different fields of view were measured from a single well. Normalization was done separately for each ligand, 100% corresponds to wells where no competitors were added and 0% corresponds to the wells where the competitors’ concentration is the largest.

**Table 3.**
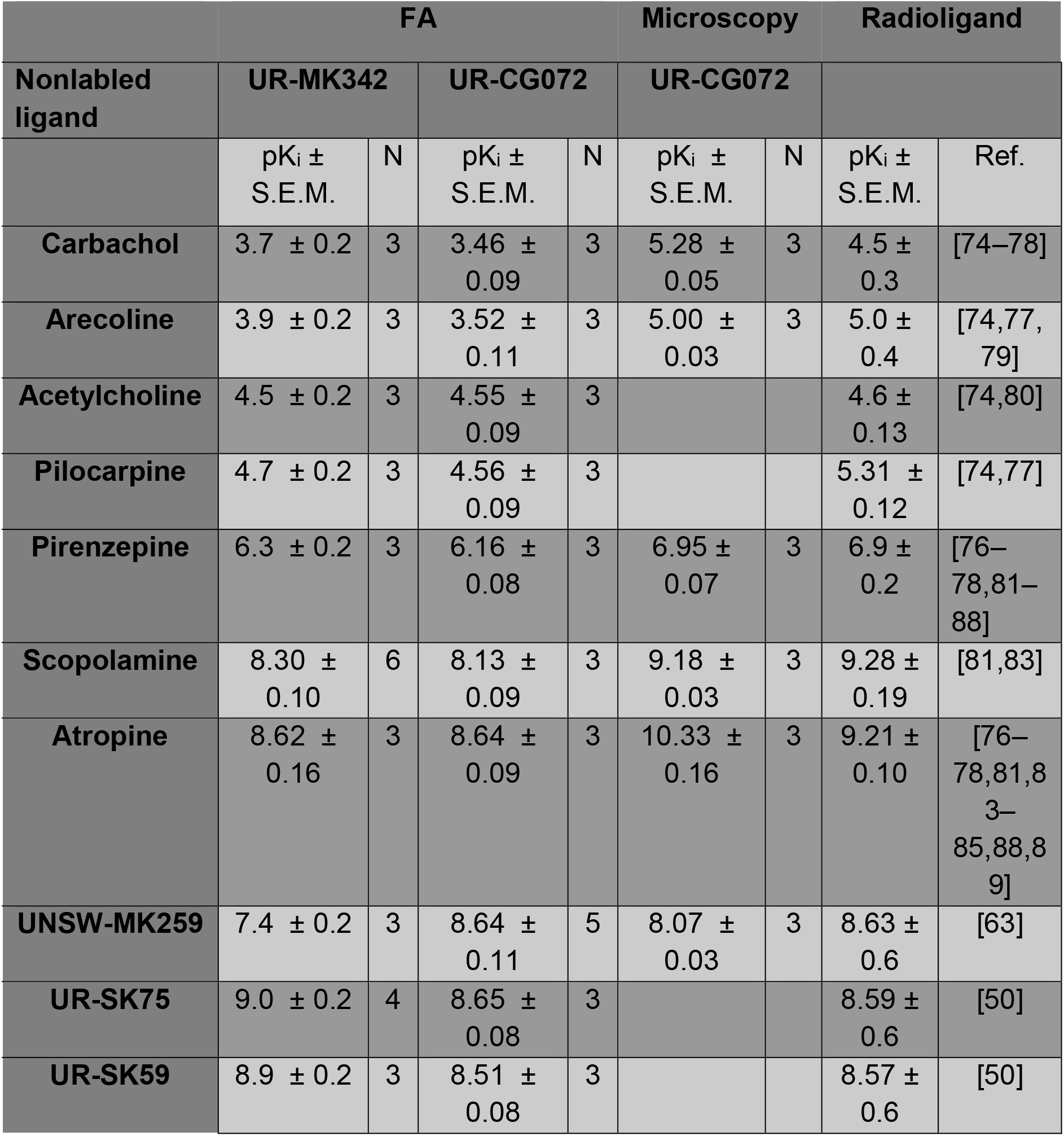
Overview of nonlabled ligand affinities

## 4. Discussion

The M_4_ receptor is connected to multiple diseases and is, therefore, an interesting target for drug development. In modern drug screening, the fluorescence-based methods have gained popularity, but the limited availability of fluorescent ligands for the M_4_ receptor has significantly hindered studies of this receptor. Recently, a set of new dibenzodiazepinone-type fluorescent ligands with high-affinity to the M_4_ receptor was synthesized, of which UR-MK342 and UR-CG072 were labeled with TAMRA [15]. In previous studies, TAMRA label has been successfully used in FA assays among other methods [90–92]. Therefore, these probes are promising candidates for developing new ligand binding assays for the M_4_ receptor. Experimental results from the FA assay show that both UR-MK342 and UR-CG072 bind to M_4_ receptors with high affinity and the K_d_ values are in good agreement with previous radioligand binding measurements. Although both ligands also have sufficiently high signals and Z-prime values to be compatible with HTS assay standards, UR-CG072 is preferred in screening assays due to its faster binding kinetics, which allows reduction of required incubation time and mitigates the effects of evaporation and potential sedimentation. Since UR-CG072 and UR-MK342 have previously been studied in the M_2_ receptor FA assay system [17], similarities and differences in both receptor systems present an opportunity to gain more insight into their binding mechanism. Interestingly, the FA value of the receptor-ligand complex remains the same regardless of which receptor subtype, M_2_ or M_4_, is measured. This similarity is evident for both fluorescent ligands. In contrast, the receptor-ligand complex FA value depends on fluorescent ligand is used and is consistently lower for UR-CG072 compared to UR-MK342 in complexes with both M_2_ and M_4_ receptor subtypes. This may indicate that the binding poses and the rotational freedom of the fluorophore moiety are similar between the two subtypes. There are also some differences in the binding properties of these probes between the FA assays of M_2_ and M_4_ receptors. Both ligands seem to show somewhat higher affinity towards the M_2_ receptor, but the differences are relatively small [17]. This is expected as orthosteric binding sites of M_2_ and M_4_ are structurally very similar [9]. However, there could be differences in the binding site accessibility since the association kinetics of both probes to the M_2_ receptor are faster compared to the M_4_ receptor. This is not surprising as ligand binding to muscarinic receptors is known to be a complex process even for somewhat smaller ligands. For example, N-methylscopolamine first binds to the allosteric site before binding to the orthosteric site of the M_2_ and M_3_ receptor subtypes [93]. The most striking difference between ligand binding to M_2_ and M_4_ receptors is the apparent lack of a clear two-phase kinetic behaviour in the case of M_4_ receptor while it is present for the M_2_ receptor. This could indicate that M_2_ and M_4_ receptor systems have differences beyond the orthosteric binding site properties as the two-phase behaviour of the M_2_ receptor-ligand binding was not specific to FA or BBV system but was also present in nanoBRET assay with mammalian cells [17]. This may indicate heterogeneity of the M_2_ receptor population, where receptors with multiple affinity states are present while the apparent heterogeneity is also ligand-dependent. For the M_4_ receptor, such heterogeneity was not observed with the ligands used in this study. The nature of this heterogeneity remains elusive but may be explained by the simultaneous existence of M_2_ receptor dimers and monomers or ligand interactions with M_2_ receptor allosteric sites. Dimerization of the M_2_ receptor is also supported by multiple previous studies while there is no information available about M_4_ receptor dimerization [94–96].

The FA method with BBV particles has many advantages such as kinetics measurement possibilities, relatively low cost, fast measurements and receptor source stability, which is achieved by using a single production batch of the BBV particle stock. Therefore, the FA based assay is a suitable option for HTS applications. However, there are also several differences between the BBV particle model system and in vivo or ex vivo conditions. For example, the live cell systems allow studying G-protein and β-Arrestin signalling and other protein-protein interactions. Furthermore, cholesterol in the membrane has an effect on ligand binding to muscarinic receptors [97], thus using live mammalian cells allows obtaining more relevant measurement results. Therefore, a live cell-based assay system, although still having notable differences from *in vivo* systems, is a significant step closer to native systems. Live-cell assays also have some general disadvantages such as slightly higher cost per experiment due to more advanced equipment required to perform the measurements and maintain cell culture. Additionally, live-cell measurements usually have higher uncertainty due to day-to-day variability. It must also be considered that the live-cell systems, which overexpress the receptors, do not fully reflect natural system and may lead to considerable biases.

The results show that receptor-ligand complex formation on the surface of live cells can be studied by automated fluorescence microscopy, which allows relatively fast measurements and high content spatio-temporal data collection. In addition, microscopy images can be used to study cell morphology, fluorophore localization, cell migration and cell death, which cannot be easily achieved with flow cytometry nor nanoBRET based measurement systems. These parameters can be useful for studying GPCR signaling [98]. Although a wide variety of advanced microscopy techniques allow measuring these parameters in great detail, high-end microscopy is often not automatic and, therefore, not suitable for high content studies. In contrast, automatic plate reader-based microscopes achieve a unique balance between the data volume and quality. This kind of automated live-cell microscopy has previously been used to study ligand binding to dopamine D_3_ receptors [35]. In the present study this method was further developed to enable the quantification of receptor-ligand binding in both equilibrium and kinetic modes. Faster kinetics of UR-CG072 compared to UR-MK342 favour using it in live-cell assays as shorter experiment times avoid negative effects such as cell detachment, changes in nutrient and oxygen concentration and cell death.

Although microscopy methods also pose some challenges related to data volumes, data analysis speed and data analysis pipeline usability, the results of this study show that suitable software and machine-learning models overcome these problems. The model comparison shows that while DL pipeline provides higher quality results the ilastik pipeline models are easier to retrain. As the final pharmacological parameters obtained with **U-Net3-BF-1** and **RF-BF-2** models are similar, with average LogIC_50_ difference of 0.15 units between models from an individual displacement curve, then both options are viable in practise depending on needed quality and useŕs level of expertise. A unique challenge with machine-learning based image analysis pipelines is the need to retrain the models if a sufficiently large domain shift is introduced into the assay such as changing the cell line or microscopy setup. Fortunately, this has to be done only once for a particular assay setup and easy to use options exist for retraining the models. Altogether, the data analysis is not a limiting factor of the proposed live-cell assay.

Using the described live-cell microscopy approach combined with machine-learning based data analysis allowed measuring ligand binding to M_4_ receptor with high quality. It is important to mention that such quality can be achieved with the fluorescence signal of bound UR-CG072 being only 4% above the background, which is substantially less than almost 200% achieved with the HEK293-D_3_R system in a previous study [35]. This lower signal is caused by a combination of multiple factors. Firstly, TAMRA fluorophore used in UR-CG072 has a lower quantum yield compared to Cy3B used in NAPS-Cy3B ligand. Secondly, the M_4_ receptor is not expressed in all CHO-K1-hM_4_R cells while the D_3_ receptor was expressed in HEK293 cells. As a final factor, using the cell culture medium instead of DPBS during imaging increases the image background intensity. Surprisingly, the reduction of absolute signal by a factor of 50 does not affect the final uncertainty of the measurements to any significant extent. The obtained R^2^ values for both saturation binding and displacement experiments are very similar for both D_3_ and M_4_ receptor microscopy assays. Essentially, it means that biological variability is the highest contributor to the total uncertainty while decreased signal has negligible uncertainty contribution. This, in turn means that this kind of assay design should work just as well with either relatively low quantum yield fluorophores or vice-versa with systems that have receptor expression more comparable to physiological expression levels if a high brightness probe is available. Thus, the proposed approach to study ligand binding to receptors has a much wider application range than previously demonstrated. Finally, the current results prove the universality of this kind of microscopy assay, as switching to another receptor and cell line did not require major changes to the analysis pipeline or assay protocol.

The developed live-cell microscopy assay can be performed in the saturation binding mode, association and dissociation kinetic modes as well as in displacement experiments for measuring the affinity of unlabeled ligands. The kinetic measurements show that the fluorescence signal is quite stable once the equilibrium is reached after the association phase. It is also evident that scopolamine induces full displacement of UR-CG072 from the M_4_ receptor as the signal reaches the same level as was in the starting point (Fig 9). Although the signal does not reach zero after dissociation, this is caused by autofluorescence, not by incomplete dissociation. UR-CG072 also has sufficiently fast kinetics for performing association and dissociation kinetics so that the morphology of CHO-K1-hM_4_R cells remains normal and the cells remain attached to the plate for the entire experiment. Both the kinetic measurements and saturation experiments prove that UR-CG072 retains its high affinity towards the M_4_ receptor in the live cell system, as expected from previous radioligand binding studies [15], while having a very low level of nonspecific binding to the cells. This makes UR-CG072 a promising fluorescent probe also for more advanced microscopy methods such as live-cell TIRF microscopy. The displacement curves obtained with UR-CG072 and unlabeled ligands have quite high quality and, therefore, this assay is suitable for the determination of affinities of unlabeled ligand binding to M_4_ receptor. The system remains stable for the duration of long experiments meaning that accurate end-point measurements can be obtained for an entire microplate even if imaging the full plate is not instantaneous. These properties also suggest that the assay can be used for small scale screening of novel ligands for example to confirm binding affinities in a live-cell system. The live-cell system is also internally consistent as the K_d_ values obtained from saturation binding measurements and kinetic measurements are in excellent agreement.

Overall, the K_d_ values of UR-CG072 obtained from both saturation and kinetic FA and live cell microscopy assays are in good agreement with each other (Table 2, Fig. 11). pK_i_ values of M_4_ receptor ligands determined with the UR-CG072 using either FA or live-cell microscopy assay, were also in good agreement (R^2^ = 0.91). The slope of the correlation was 0.84 while the intercept was 2.2. The live cell method systematically estimates higher affinities for low affinity ligands while for high-affinity ligands in the nanomolar range the estimated values are numerically more similar between the assays (Fig. 11). However, most low affinity ligands are agonists while high-affinity ligands are antagonists. Therefore, it is difficult to determine whether there is a systematic difference between assays for low affinity ligands or simply agonists. Agonism causing the systematic difference is theoretically well founded, as the high-affinity receptor state is usually stabilized by G-proteins, which are not present in the BBV particles. A similarly good correlation was previously found between nanoBRET assay and FA assay using the same probe with M_2_ receptor (R^2^ = 0.94) with the same systematic differences between the pK_i_ values measured in BBV particles and live-cells [17]. This further supports that the systematic difference between the determined agonist pK_i_ values is caused by differences between BBV particle and live-cell systems.

**Fig. 11.**
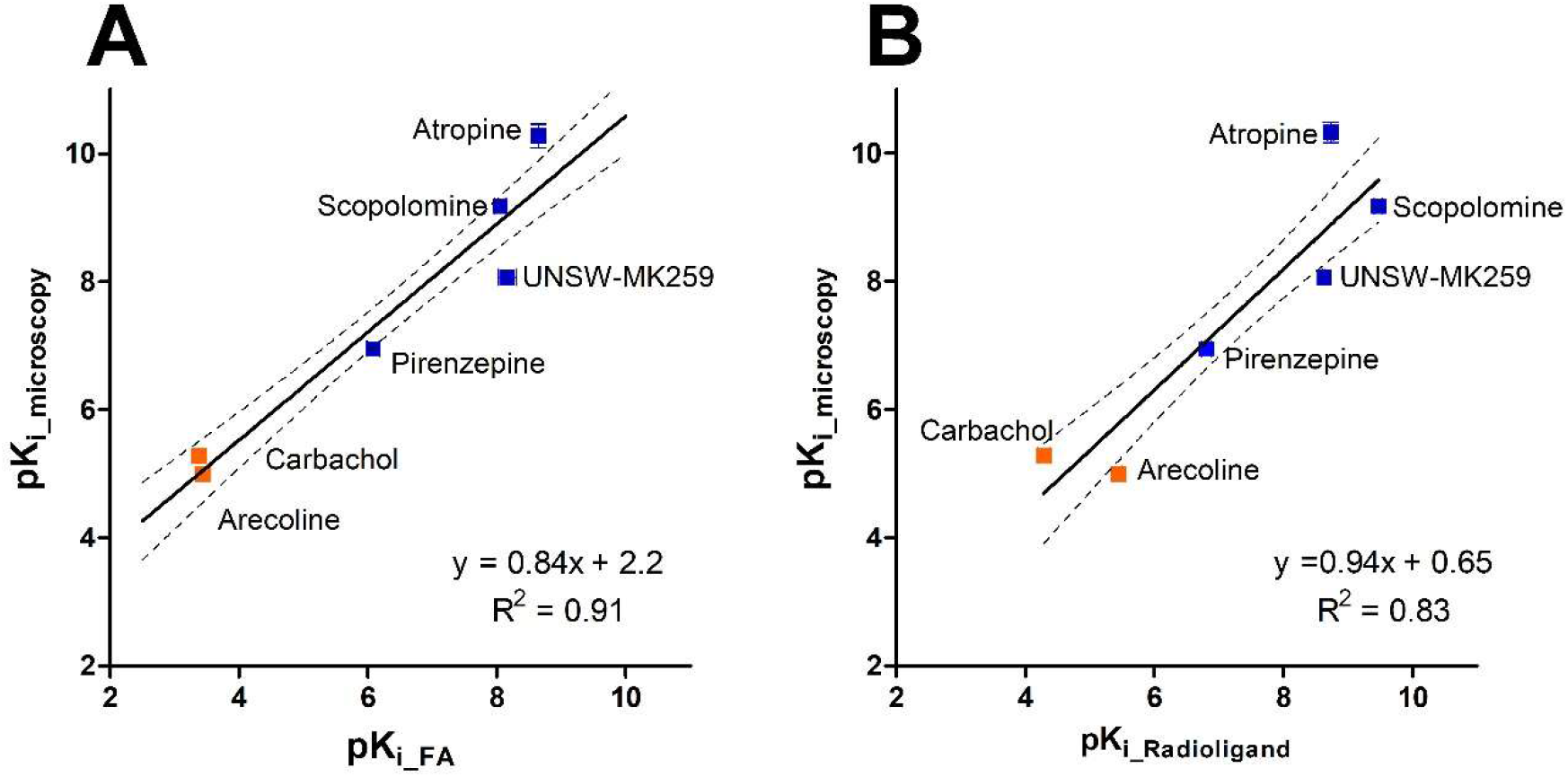
Correlation plots of affinities (pK_i_ values) of reported muscarinic receptor ligands measured with UR-CG072 in different assay systems. (A) Comparison of pK_i_ values determined in the microscopy competition binding assay and pK_i_ values obtained from the FA competition binding assay. (B) Comparison of pK_i_ values determined in the microscopy competition binding assay and pK_i_ values obtained from radioligand binding assay from literature (Table 3). Investigated agonists are presented as orange symbols 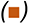, antagonists as blue symbols 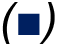. Black lines represent the best linear relationship between the datasets and the dashed black line represent 95% confidence bands.

The developed live cell microscopy assay can be modified for wider applications in the future. One development direction is further assay automatization by removing the remaining manual steps from the data analysis process. This could also include an even more standardized pipeline for machine learning model development or a larger set of pretrained models that cover the detection of most common cell lines. We believe that it can be further developed to an extent at which the live-cell microscopy could also be used in an HTS context. Another development direction is a shift towards more natural systems such as tissue preparations, live tissues or tumor spheroids and measuring additional downstream signaling events in addition to ligand binding. Using these more challenging systems requires finding suitable fluorophores to overcome tissue autofluorescence and ligands with suitable kinetic properties to slow down fluorescence ligand dissociation during washing steps. Additionally, more advanced DL models may be necessary. The present study serves as a solid foundation for such developments.

As for the more general unlabeled ligand screening, both FA and live-cell microscopy methods and fluorescence ligands could be utilized as a rapid and convenient options for guiding the synthesis of novel M_4_ receptor ligands and allosteric modulators. Both methods also allow for kinetic measurements which may help uncover more detailed binding mechanisms. Overall, choosing the suitable method for a specific experiment highly depends on the required throughput and availability of equipment. While FA method with BBV particles fulfils many requirements of HTS applications, live-cells are a vastly more flexible option for studying complex signaling pathways. Therefore, live-cell microscopy-based ligand binding assays are likely to have an ever-growing role in the future of ligand binding studies.

## CRediT authorship contribution statement

**Maris-Johanna Tahk**: Writing - Original Draft, Investigation, Formal analysis, Visualization, Methodology, Data Curation, Validation; **Jane Torp**: Writing Original Draft, Investigation, Software, Formal analysis, Visualization, Methodology, Validation Data Curation,; **Mohammed A.S. Ali:** Software, Visualization, Writing - Review & Editing; **Dmytro Fishman:** Resources, Writing - Review & Editing, Supervision; **Leopold Parts**: Supervision, Resources, Funding acquisition, Conceptualization, Writing - Review & Editing; **Lukas Grätz**: Resources, Writing - Review & Editing; **Christoph Müller**: Resources Writing - Review & Editing; **Max Keller**: Resources, Conceptualization Writing - Review & Editing, Funding acquisition; **Santa Veiksina:** Funding acquisition, Resources, Writing - Review & Editing; **Tõnis Laasfeld**: Investigation, Software, Writing - Original Draft, Conceptualization, Formal analysis, Visualization, Methodology, Data Curation; **Ago Rinken**: Conceptualization, Funding acquisition, Supervision, Writing - Original Draft, Resources

## Acknowledgements

This publication was supported by the University of Tartu ASTRA Project PER ASPERA, financed by the European Regional Development Fund. This study was supported by the Estonian Research Council grant (PSG230), by the COST action CA 18133 ERNEST, by the Research Training Group GRK1910 of the Deutsche Forschungsgemeinschaft (DFG), Wellcome (206194), and the Estonian Centre of Excellence in IT (EXCITE) (TK148).

We would like to thank Dr. Anni Allikalt for cloning the M_4_ receptor into the pFastBac vector.

## Abbreviations

5-TAMRA: 5-Carboxytetramethylrhodamine
BBV: budded baculovirus
BRET: bioluminescence resonance energy transfer
CHO-K1: Chinese hamster ovary cells
DiI: 1,1’-Dioctadecyl-3,3,3’,3’-Tetramethylindocarbocyanine Perchlorate
DL: deep learning
DMEM/F12: Dulbecco’s Modified Eagle Medium/Nutrient Mixture F-12
DPBS: Dulbecco’s phosphate-buffered saline
FA: fluorescence anisotropy
GPCR: G-protein-coupled receptors
GUI: graphical user interface
HEK293-D3R: Human embryonic kidney 293 cells expressing dopamine D3 receptors
HTS: high-throughput screening
hM4: human muscarinic acetylcholine receptor M4 subtype
IC: Intracellular area
NMBG: near-membrane background
mAChR: muscarinic acetylcholine receptor
MB: membrane
MCC: Matthew’s correlation coefficient
MOI: multiplicity of infection
RF: random forest
U-Net3-FL-1: U-Net3 architecture based fluorescence image cell segmentation model
RF-FL-1: Random Forest based fluorescence image cell segmentation model
U-Net3-BF-1: U-Net3 architecture based bright-field image cell segmentation model
RF-BF-1: Random Forest based bright-field image cell segmentation model version 1
RF-BF-2: Random Forest based bright-field image cell segmentation model version 2

